# Familial Alzheimer disease mutation identifies novel role of SORLA in release of neurotrophic exosomes

**DOI:** 10.1101/2025.04.22.649924

**Authors:** Kristian Juul-Madsen, Ina-Maria Rudolph, Jemila P. Gomes, Katrina Meyer, Peter L. Ovesen, Malgorzata J. Górniak-Walas, Marianna Kokoli, Narasimha S. Telugu, Malthe von Tangen Sivertsen, Duncan S. Sutherland, Johan Palmfeldt, Sebastian Diecke, Olav M. Andersen, Matthias Selbach, Thomas E. Willnow

## Abstract

Sortilin-related receptor with A-type repeats (SORLA) is an intracellular sorting receptor that directs target proteins between endocytic and secretory compartments of cells. Mutations in *SORL1*, encoding SORLA, are common in individuals suffering from Alzheimer disease (AD) of unknown etiology. Conceptually, characterization of inheritable *SORL1* variants associated with AD can provide important new information about functions of this receptor relevant for aging brain health. Here, we focused on elucidation of the AD-associated variant SORLA ^N1358S^, carrying a mutation in the main ligand binding domain of the receptor. Using unbiased quantitative proteome screens, we identified major alterations in the mutant receptor interactome linked to biogenesis and secretion of exosomes. Using advanced biophysical, cell biological, as well as functional studies in stem cell-derived human cell models we corroborated impaired release and loss of neurotrophic action of exosomes from neurons and microglia expressing SORLA^N1358S^. An impaired neurotrophic potential was attributed to an altered exosomal content of RNA binding proteins and associated microRNAs, known to control neuronal growth and maturation. Our studies identified a so far unknown function for SORLA in controlling the quantity and trophic quality of extracellular vesicles secreted by cells, and they argue for impaired cellular cross talk through exosomes as a pathological trail contributing to the risk of AD seen with carriers of *SORL1* variants.

## INTRODUCTION

Sortilin-related receptor with A-type repeats (SORLA) is a 230 kDa type-1 transmembrane protein expressed in various mammalian cell types, including neurons and microglia in the human brain (reviewed in ^1,2^). SORLA acts as an intracellular sorting receptor directing multiple target proteins between Golgi, cell surface, and endo-lysosomal compartments, sorting paths central to endocytic and secretory functions of cells. SORLA is best known for its ability to act as neuronal sorting receptor for the amyloid precursor protein (APP), preventing its proteolytic breakdown into amyloid-β peptides (Aβ), a causative agent in Alzheimer disease (AD) ^3–6^. Intriguingly, genetic studies have identified the encoding gene *SORL1* as a novel disease gene in familial forms of AD (FAD). In fact, *SORL1* variants predicted *in silico* as damaging to protein structure may be present in as many as 3% of all patients with FAD of unknow etiology ^7–9^. This prediction is supported by studies in cultured cells, documenting impaired folding and maturation of the receptor polypeptide seen with many *SORL1* gene variants ^10^. A second class of *SORL1* variants, associated with AD thus far, exhibit disrupted intracellular sorting of the receptor and its target APP, corroborating control of amyloidogenic processing of APP as a receptor function of disease relevance ^10–13^. Still, whether association with FAD is solely explained by the ability of SORLA to sort APP in neurons, or whether yet unknown receptor functions contribute to the risk of the disease seen with some *SORL1* alleles, remains an open question. In line with this notion, prior analysis of mutation G511R in the extracellular domain of the receptor identified binding of Aβ to this domain as another function of SORLA lost in FAD ^14^.

Conceptually, the unbiased study of loss-of-function variants in *SORL1* associated with FAD may enable us to uncover novel receptor interactions crucial to aging brain health not constrained by prior hypotheses of receptor actions. Here, we focused on the functional characterization of the mutation N1358S as it localizes to the complement-type repeats, a major ligand binding domain in the receptor polypeptide. Combining unbiased interactome studies with targeted analyses of iPSC-derived human brain cell types, we identified impaired biogenesis and loss of neurotrophic actions of exosomes as cellular phenotypes caused by expression of SORLA^N1358S^. These findings document a novel role for SORLA in cell-to-cell communication through exosomes, and suggest defects in these processes to contribute to AD pathology in carriers of the *SORL1^N1358S^*variant.

## RESULTS

### Mutation N1358S alters the SORLA interactome

To investigate the effects of inheritable *SORL1* variants on SORLA activities potentially relevant to AD, we focused on mutation N1358S identified in an individual with early onset AD ^7,8^. Our choice was guided by the fact that this mutation localizes to the cluster of complement-type repeats (CR), a major ligand binding domain in the receptor polypeptide (Fig. 1a) ^15,16^. Specifically, N1358S localizes to CR number 7 (CR7) and is predicted to have little effect on the overall conformation of this CR or adjacent receptor domains (Fig. 1b). From analysis of this SORLA variant, we hoped to elucidate novel mechanisms whereby altered ligand interactions of this multifunctional receptor may explain its (path)physiological roles in brain health and disease.

**Figure 1:**
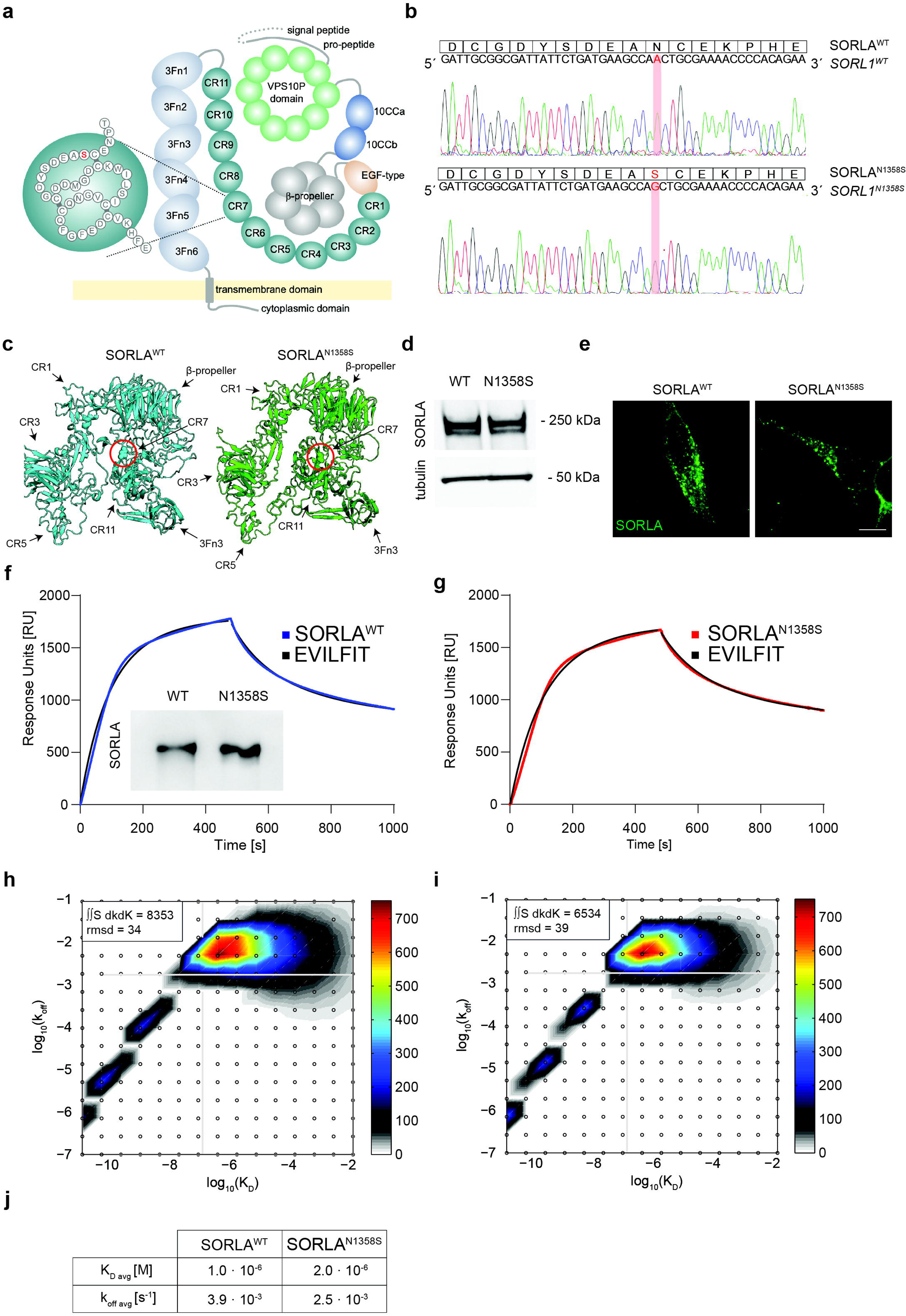
Mutation N1358S does not impact APP binding to SORLA. (**a**) Structure of SORLA highlighting mutation N1358S in complement-type repeat (CR) 7. EGF, epidermal growth factor repeat; 3Fn, fibronectin type 3 domain. (**b**) Alphafold prediction of the impact of N1358S in CR7 on SORLA structure (amino acids K497 to R1883). N1358S is highlighted using spheres. (**c**) Sequence of expression constructs for SORLA^WT^ and SORLA^N1358S^ introduced into cell line SY5Y. Corresponding peptide sequences are shown above. (**d**) SORLA levels in lysates of SY5Y cell lines expressing SORLA^WT^ or SORLA^N1358S^. Detection of tubulin served as loading control. Migration of molecular weight markers (in kDa) in the gel are indicated. (**e**) Immunofluorescence detection of SORLA (green) in SORLA^WT^ and SORLA^N1358S^ SY5Y cell lines. Scale bar: 20 µm. (**f**, **g**) Surface plasmon resonance (SPR) analysis of binding of purified SORLA^WT^ (f) and SORLA^N1358S^ ectodomains (g) to the ectodomain of APP^695^ coupled to the sensor chip surface. Sensorgrams for SORLA variants binding to APP are given as blue (f) or red (g) lines. EVILFIT is shown as black lines in both graphs. Western blot analysis (inset in F) documents integrity of purified SORLA ectodomains used for SPR analysis. **(h, i)** EVILFIT distributions presented as contour plots on 2D grids with log10(K_D_) and log10(k_off_) on the x- and y-axes, respectively, at signal max 750 response units (RU) for SORLA^WT^ (h) and SORLA^N1358S^ (i). The root-mean square deviation (rmsd) between the experimental data and the model is stated for each sensorgram in the top left corner. (**j**) K_D_ and k_off_ values for SORLA variants binding to APP^695^ derived from integration of EVILFIT.

Initially, we used a PCR-based mutagenesis strategy to generate expression constructs encoding human wildtype (SORLA^WT^) and mutant (SORLA^N1358S^) receptor variants (Fig. 1c). When stably expressed in the neuroblastoma cell line SY5Y, both receptor variants showed similar protein levels (Fig. 1d) as well as localization to intracellular vesicular compartments (Fig. 1e), confirming that mutation N1358S does not overtly impact biosynthesis and cellular localization of SORLA. As CR domains have been identified as the binding site for APP before ^15,16^, we initially queried whether N1358S may impair interaction with SORLA^N1358S^ using surface plasmon resonance (SPR) analyses. To do so, we expressed hexa-His-tagged versions of the soluble wildtype and mutant SORLA ectodomains in HEK293 cells (inset in Fig. 1f) and compared their abilities to bind the ectodomain of APP^695^ coupled to the sensor chip surface. These studies revealed comparable binding kinetics at neutral pH with SPR sensorgram peaks at 1750 and 1650 response units (RU) for SORLA^WT^ and SORLA^N1358S^, respectively (Fig. 1f-g).

For analysis of SPR data, the Langmuir model is typically used to describe protein interactions from saturation of surface binding sites at steady-state concentration ^17^. However, this model only considers the possibility of a 1:1 interaction between ligand and analyte. To interrogate whether the interaction of SORLA with APP may in fact involve several independent CR sites, only one of which may be affected by N1358S, we re-analyzed the SPR data using EVILFIT. This algorithm permits the description of SPR data as an ensemble of multiple pseudo-first-order reactions (see supplementary methods for details) ^18^. The resulting 2D distribution of minimally required 1:1 interactions to model the data is plotted in a grid of resulting K_D_ and k_off_ rates (Fig. 1h and i). EVILFIT was in perfect agreement with the SPR analyses (black graphs in Fig. 1f and g). It confirmed that the binding of both SORLA^WT^ and SORLA^N1358S^ to APP^695^ was dominated by a single type of interaction (shown in the upper right corner of the 2D grid in Fig. 1h and i), with no obvious differences in K_D_ and k_off_ rates seen between the two receptor variants (Fig. 1j).

As the binding between SORLA and APP was not affected by N1358S, we considered that this mutation in the ligand binding domain may alter interaction with yet unknown receptor targets. Therefore, we took an unbiased proteomics approach, not constrained by prior hypotheses, to compare the interactomes of SORLA^WT^ and SORLA^N1358S^ in cells. In detail, we applied stable isotope labeling by amino acids in cell culture (SILAC) to metabolically label SY5Y cells expressing SORLA^WT^ or SORLA^N1358S^ with heavy or light isotopes of lysine and arginine. Subsequently, SORLA interacting proteins were precipitated from the cell lysates using anti-SORLA IgG-coupled beads and subjected to mass spectrometry-based proteomics (Fig. 2a). Isotope labelling did not differentially impact input levels of the cell proteome as exemplified by comparing input intensities for SORLA^WT^ cells labeled with either heavy or light isotopes (Fig. 2b). Comparing the SY5Y^WT^ interactome enriched with anti-SORLA versus non-IgG-coupled beads mainly documented hits in the anti-SORLA immunoprecipitate (IP), including the receptor and its ligand APP (Fig. 2c, upper right corner). By contrast, very few proteins unique to the non-IgG IP were detected (Fig. 2c, lower left corner), demonstrating the specificity of our assay for identifying proteins directly or indirectly interacting with SORLA.

**Figure 2:**
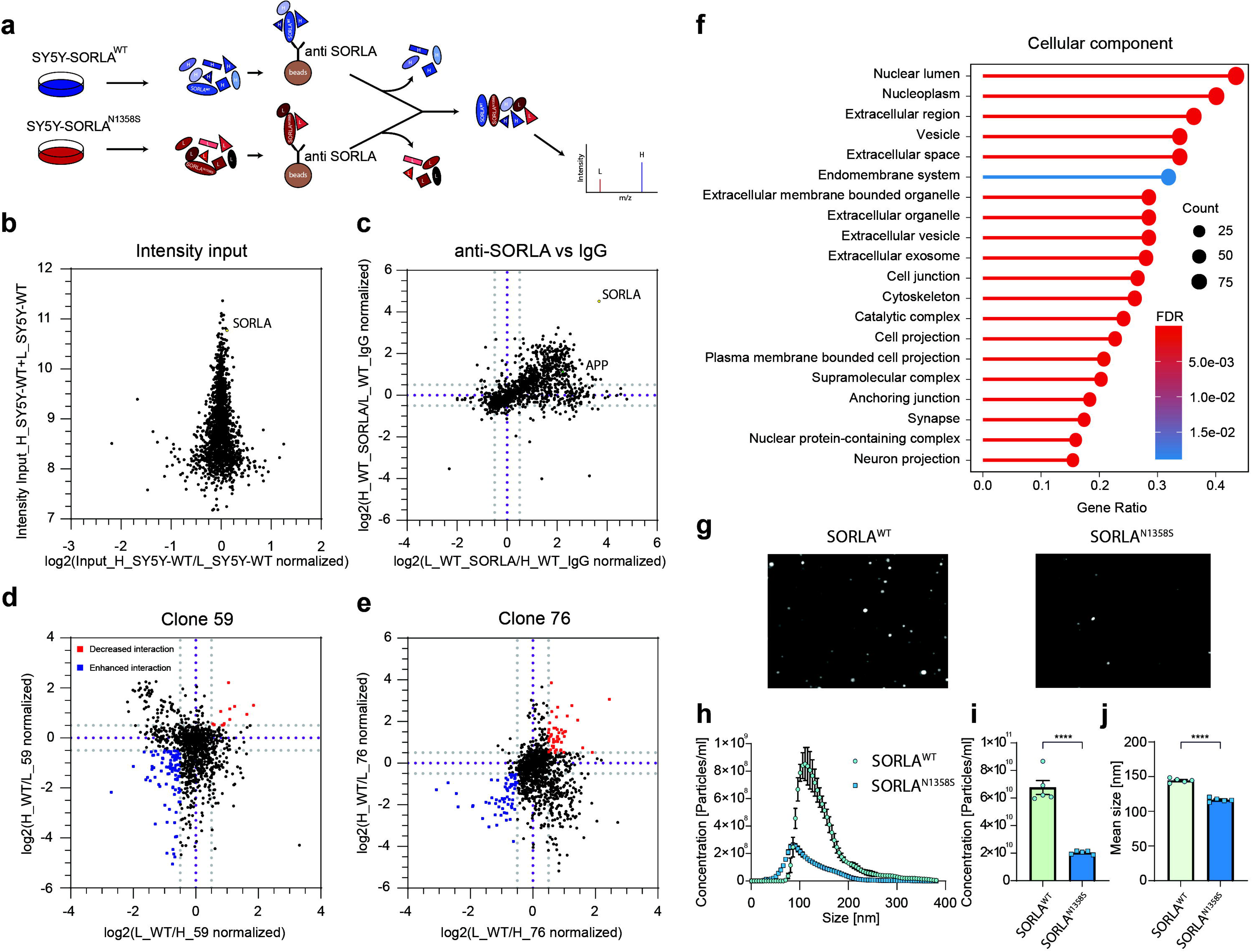
Mutation N1358S alters the interactome for SORLA involved in exosome biogenesis. (**a**) Workflow for stable isotope labeling by amino acids in cell culture (SILAC). SY5Y cell lines stably expressing SORLA^WT^ or SORLA^N1358S^ were labeled with heavy (H) or light (L) isotopes. Subsequently, cell lysates were immunoprecipitated using anti-SORLA or non-immune (negative control) IgG coupled to beads. Immunoprecipitates from both cell lines were combined and subjected to mass spectrometry-based proteomics. (**b**) Relative intensity inputs of proteins extracted from SY5Y^WT^ cells labelled with heavy or light isotopes are given, documenting that neither heavy nor light isotope labeling has noticeable effects on input. The signal representing SORLA is indicated. (**c**) Comparison of interacting proteins immunoprecipitated with anti-SORLA as compared to unspecific IgG from SY5Y^WT^ cells. Purple lines indicate no difference in interaction, grey lines indicate >50 % change in interaction. Most protein hits precipitate with anti-SORLA beads (upper right corner above dashed grey lines) but not with unspecific IgG beads (lower left corner below grey dashed lines). Signals representing SORLA and APP in the anti-SORLA precipitate are indicated. (**d**, **e**) Analysis of interacting proteins precipitated with anti-SORLA IgG beads from SY5Y^WT^ or SY5Y^N1358S^ cell clones 59 (d) and 76 (e). Proteins interacting relatively stronger with SORLA^N1358S^ are given as blue dots (>50% enriched as compared to SORLA^WT^). Hits with reduced mutant receptor interaction are shown as red dots (>50% reduced compared to SORLA^WT^). (**f**) Over-representation analysis showing altered interaction with SORLA^N1358S^ (clones 59 and 76) as compared to SORLA^WT^. Gene ontology (GO) analysis of over-represented proteins in the SORLA interactome related to the cellular component pathways (visualized with Genekitr). Benjamini-Hochberg test was applied to calculate adjusted p values of GO terms. (**g**) Representative images of purified exosomes analyzed by nanoparticle tracking (NTA, scatter detection mode) document a reduced number of particles released by SY5Y^N1358S^ as compared to SY5Y^WT^ cells. (**h-j**) Size distribution (h), total concentration (i), and mean size (j) of exosomes secreted from SY5Y^WT^ and SY5Y^N1358S^ as determined by NTA. SY5Y cells expressing SORLA^N1358S^ release fewer and also smaller exosomes as compared to cells expressing SORLA^WT^. Data represent five independent replicates analyzed using Student’s t-test (p < 0.0001).

Following validation, we applied our SILAC protocol to input from SY5Y^WT^ cells and two SY5Y cell clones expressing SORLA^N1358S^ (clones 59 and 76). To control for isotope labeling effects, all three cell lines were individually labeled with either heavy or light isotopes. Our studies identified a significant shift in receptor interactions, with multiple hits showing enhanced or decreased interaction with the mutant receptor (Figs. 2d and e). When analyzed by over-representation analysis, proteins differentially interacting with SORLA^N1358S^ showed enrichment for cellular components related to extracellular vesicle biogenesis and release. Specifically, the top 20 most over-represented cellular pathways included the terms vesicle, extracellular space, extracellular membrane-bound organelle, extracellular vesicle, as well as exosome (Fig. 2f, Suppl. table S1).

### SORLA^N1358S^ impairs the release of exosomes from SY5Y cells

Given the predominance of pathways related to extracellular vesicle biogenesis over-represented in the differential analysis of SORLA^WT^ versus SORLA^N1358S^ interactomes, we focused on a possible role for the receptor in this biological process not recognized so far. Specifically, we considered the relevance of SORLA for biosynthesis of exosomes, a process potentially linked to its presumed function in endo-lysosomal sorting processes ^19–21^. To test our hypothesis, we purified exosomes from the supernatant of SY5Y cells expressing SORLA^WT^ or SORLA^N1358S^ using established protocols of differential ultracentrifugation. These protocols enrich for exosomes with sizes between 50-200 nm ^22^. Following purification, we applied nanoparticle tracking analysis (NTA) to compare the number and size of particles released by both cell lines. NTA tracks light scattering by single particles to determine their Brownian motion in suspension and to derive their amount and size thereof ^23^. In these analyses, a significant decrease in the amounts of exosomes released by SY5Y^N1358S^ as compared to SY5Y^WT^ cells was evident (Fig. 2g). Detailed analyses of particle size distribution and quantity (Fig. 2h) documented a 70% reduction in concentration (Fig. 2i) and a 20% reduction in average size (Fig. 2j) for exosomes released by SY5Y^N1358S^ as compared to exosomes from SY5Y^WT^. These findings provided a functional correlate to the results of the interactome studies in cell free systems using SILAC, and confirmed a prospective role for SORLA in exosome release from neuroblastoma cells.

### Expression of SORLA^N1358S^ reduces the amount exosomes released by human neurons and microglia

To query whether SORLA modulates the biogenesis of exosomes by more physiologically relevant human brain cell types, we used CRISPR/Cas9-based genome editing to generated isogenic human iPSC lines, either wildtype (SORLA^WT^) or genetically deficient for SORLA (SORLA^KO^), or homozygous for SORLA^N1358S^ (Suppl. figure 1a-b). Successful ablation of SORLA expression in KO iPSC and comparable expression levels of SORLA^WT^ and SORLA^N1358S^ in this cell type was confirmed by Western blotting (Suppl. figure 1c). SORLA deficiency, or mutant receptor expression, did not impact iPSC pluripotency as tested by expression of pluripotency markers using quantitative (q) RT-PCR (Suppl. figure S1e-f) and immunocytochemistry (Suppl. figure 1g).

Using standard protocols of forced overexpression of transcription factor NGN2 ^24^, we differentiated our three iPSC lines into induced cortical neurons (iN; Fig. 3a-b) with no apparent impact of *SORL1* genotypes on the appearance of neurons (Fig. 3b) or expression of neuronal marker micro-tubule associated protein 2 *(MAP2*) (Fig. 3c). To test the impact of *SORL1* genotypes on exosomal biogenesis, we purified exosomes from the supernatants of all three iN populations and subjected them to NTA (Fig. 3d). While the size of purified vesicles was comparable between the genotypes (Fig. 3e and g), a clear reduction in the concentration of exosomes released by SORLA^KO^ and SORLA^N1358S^ iN was evident when compared to SORLA^WT^ iN (Figs. 3e and f).

**Figure 3:**
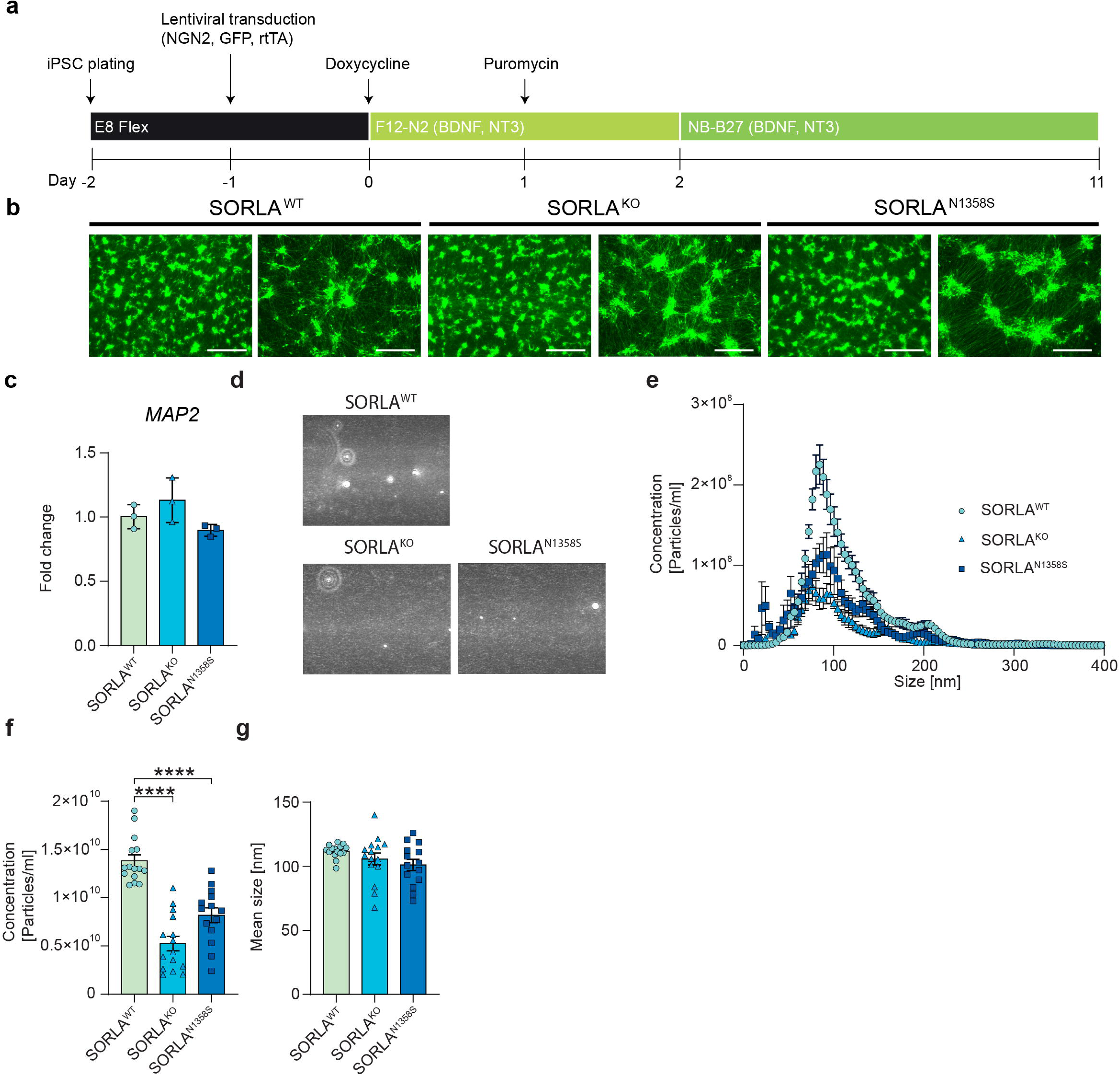
Loss of SORLA, or expression of SORLA^N1358S^, decreases the amount of exosomes released by human cortical neurons. (**a**) Differentiation protocol used to generate induced cortical neurons (iNs) using lentiviral expression constructs for NGNG2, rTA, and GFP ^24^. See supplementary methods for details. (**b**) Fluorescence detection of GFP for SORLA^WT^, SORLA^KO^ and SORLA^N1358S^ iN at day 11 of differentiation. Scale bars: 750 μm (right images), 300 μm (left images). (**c**) Transcript levels of neuronal marker *MAP2* in the indicated iN lines at day 11 of differentiation. Relative quantifications (RQ) of fold change represent 2-ddCt relative to iPSC with *GAPDH* used as reference gene. (**d**) Exosomes purified from SORLA^WT^, SORLA^KO^, or SORLA^N1358S^ iN were analyzed using scatter detection mode of nanoparticle tracking analysis (NTA). The numbers of particles in SORLA^KO^ and SORLA^N1358S^ are lower as compared to SORLA^WT^ iN. (**e-g**) NTA analysis of size distribution (e), total concentration (f), and mean size (g) of exosomes (50 – 200 nm size rage) secreted by SORLA^WT^, SORLA^KO^, and SORLA^N1358S^ iN. Cells lacking SORLA or expressing the mutant receptor release significantly fewer exosomes as compared to cells expressing the wildtype receptor (e, f). No difference is seen in exosome size comparing genotypes (e, g). Data in c-e are derived from 3 independent differentiations and exosome purifications per genotype, with 5 technical replicates each. Data were analyzed using one-way ANOVA with Tukey’s correction for multiple testing (****, p < 0.0001).

Conceptually, a role for SORLA in exosomal biogenesis may be a function unique to neurons or represent an activity common to other (brain) cells. Given the emerging role of SORLA action in microglia as a possible contributor to AD pathology ^19,25–27^, we tested whether SORLA may also control the release of exosomes from this cell type. Thus, all three iPSC lines were differentiated into induced human microglia (iMG) using published protocols ^28^. Differentiation produced cells with ramified microglia-like morphology (Fig. 4a, day 38) with no discernable difference in expression of microglia markers *IBA1*, *P2RY12*, or *ITGAM* comparing genotypes by qPCR analyses (Fig. 4b). Immunostainings confirmed robust expression of SORLA in SORLA^WT^ and SORLA^N1358S^ iMG, but absence from SORLA^KO^ iMG (Fig. 4c). Electron microscopical inspection of purified extracellular vesicles released by iMG of all three genotypes identified vesicles of sizes ranging between 50-200 nm (Fig. 4d). In line with our preparation protocol ^29^, a fraction of these vesicles showed a cup-like shape, representative of exosomes (Fig. 4d). While no apparent difference in structure of exosomes was noted comparing genotypes, their concentrations in iMG supernatants were significantly different as documented by NTA. As shown in Fig. 4f-h, the concentration of vesicles released by iMG expressing the wildtype receptor was significantly higher than for exosomes released by cells expressing SORLA^N1358S^ or lacking the receptor.

**Figure 4:**
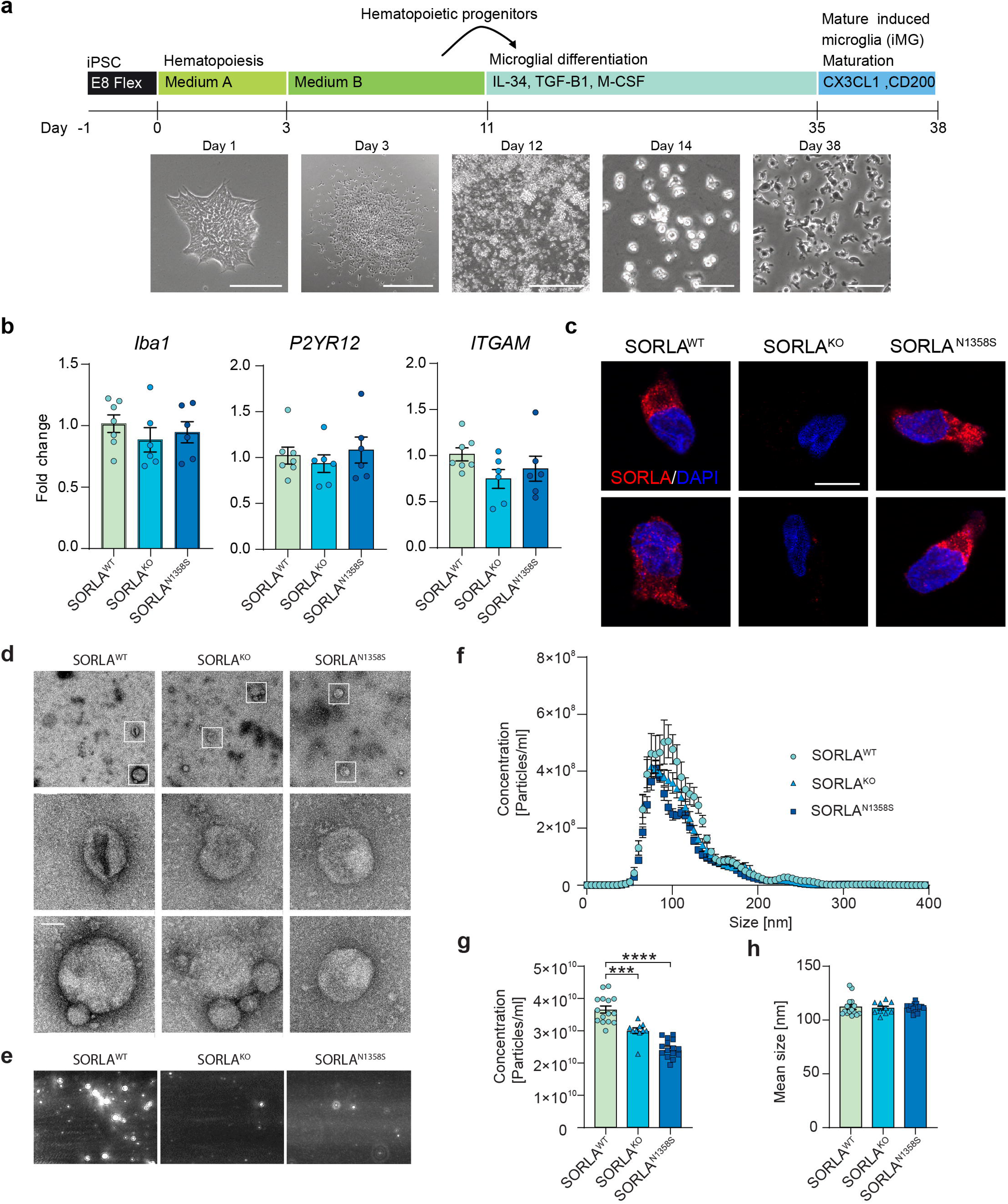
Loss of SORLA, or expression of SORLA^N1358S^, impaires the amount of exosomes released by human microglia. (**a**) Differentiation protocol used to generate induced human microglia (iMG). Phase contrast images of iPSCs, hematopoietic progenitors (HPs), and iMG at different stages of differentiation are shown below. Scale bars: 1000 μm (day 3 and 12), 200 μm (day 38). (**b**) Transcript levels of microglia markers *IBA1*, *P2YR12*, and *ITGAM* in the indicated iMG lines (day 38). Relative quantifications (RQ) of fold change represent 2-ddCt relative to iPSC, with *GAPDH* and *TBP* used as reference genes. (**c**) Immunofluorescence detection of SORLA (red) in SORLA^WT^, SORLA^KO^, and SORLA^N1358S^ iMG. Nuclei were counterstained with DAPI (blue). Scale bar: 10 μm. **(d)** Electron microscopical analysis of exosomes purified from the supernatants of iMG expressing SORLA^WT^ or SORLA^N1358S^, or being genetically deficient for the receptor (SORLA^KO)^. Images are representative of three independent exosome collections per genotype. White boxes (in overview) indicate exosomes shown in the higher magnification images below. Scale bars: 200 nm (upper row) or 50 nm (lower and middle rows). (**e**) Exosomes purified from SORLA^WT^, SORLA^KO^, or SORLA^N1358S^ iMG were analyzed using scatter detection mode of nanoparticle tracking analysis (NTA). The number of particles is comparable in SORLA^KO^ and SORLA^N1358S^, but lower when compared to SORLA^WT^ iMG. (**f-h**) NTA analyses of size distribution (f), total concentration (g), and mean size (h) of exosomes secreted by SORLA^WT^, SORLA^KO^, or SORLA^N1358S^ iMG. Cells lacking SORLA or expressing the mutant receptor release significantly fewer exosomes as compared to cells expressing the wildtype receptor (f, g). No difference is seen in exosome size comparing cell types (f, h). Data in d-h are derived from 2-3 independent differentiations and exosome purifications per genotype, with 5 technical replicates each. Data were analyzed using one-way ANOVA with Tukey’s correction for multiple testing (****, p < 0.0001).

Exosome biogenesis is a complex process, intimately linked to endocytic vesicle trafficking (reviewed in ^30,31^). To test spatial proximity of SORLA to cellular compartments in exosome biogenesis, we studied its colocalization with three established markers of exosome formation, namely members of the tetraspanin superfamily CD9, CD63, and CD81 ^32^. In iMG, SORLA^WT^ and SORLA^N1358S^ both colocalized with all three exosomal biogenesis markers to a comparable extent (Suppl. figure 2a-c), although a subtle difference in the Pearson’s correlation coefficient was noted for CD63 (Suppl. figure 2d). Interestingly, the total amount of intracellular immunosignals for CD9 and CD81 were significantly increased in SORLA^N1358S^ as compared to SORLA^WT^ iMG, corroborating an exosome release defect in this genotype (Suppl. figure 2e).

Taken together, our data supported a role for wildtype SORLA in exosome biosynthesis pathways, an activity relevant for neuronal and microglial cell types in the human brain.

### Expression of SORLA^N1358S^ impairs the composition and trophic action of exosomes released by human microglia

Among other functions, microglia-derived EVs have been associated with support of neuronal maturation, such as neurite outgrowth ^33–35^. As well as by quantity, exosomes released from SORLA^KO^ or SORLA^N1358S^ iMG were also distinguished by impaired neurotrophic quality from SORLA^WT^ vesicles. This fact was documented when we tested the ability of purified exosomes to promote neurite growth and branching in immature neurons. This assay is commonly used to score functional properties of exosomes in promoting neuronal maturation ^34,35^.

In our test, we added iMG-derived exosomes to human iPSC induced to differentiate into neurons by stably overexpression of the transcription factor NGN2 ^36^. These neurons were wildtype for the *SORL1* gene. Identical amounts of purified exosomes (10^5^ exosomes/ml) were added to replicate layers of neurons to adjust for the different quantities of vesicles released by SORLA^KO^ and SORLA^N1358S^ as compared to SORLA^WT^ microglia. Neurite growth and branching was scored in exosome-treated or untreated immature neurons for 12 h using the IncuCyte live imaging system and the build-in neuro-track software package (Fig. 5a). In these experiments, exosomes from SORLA^WT^ iMGs promoted neurite growth (Fig. 5b and d) and branching (Fig. 5c and e) when compared with untreated neurons, documenting their neurotrophic potential. By contrast, exosomes purified from the supernatants of iMG either SORLA^N1358S^ or SORLA^KO^ failed to promote neurite growth (Fig. 5b and d) and branching (Fig. 5c and e) over untreated controls.

**Figure 5:**
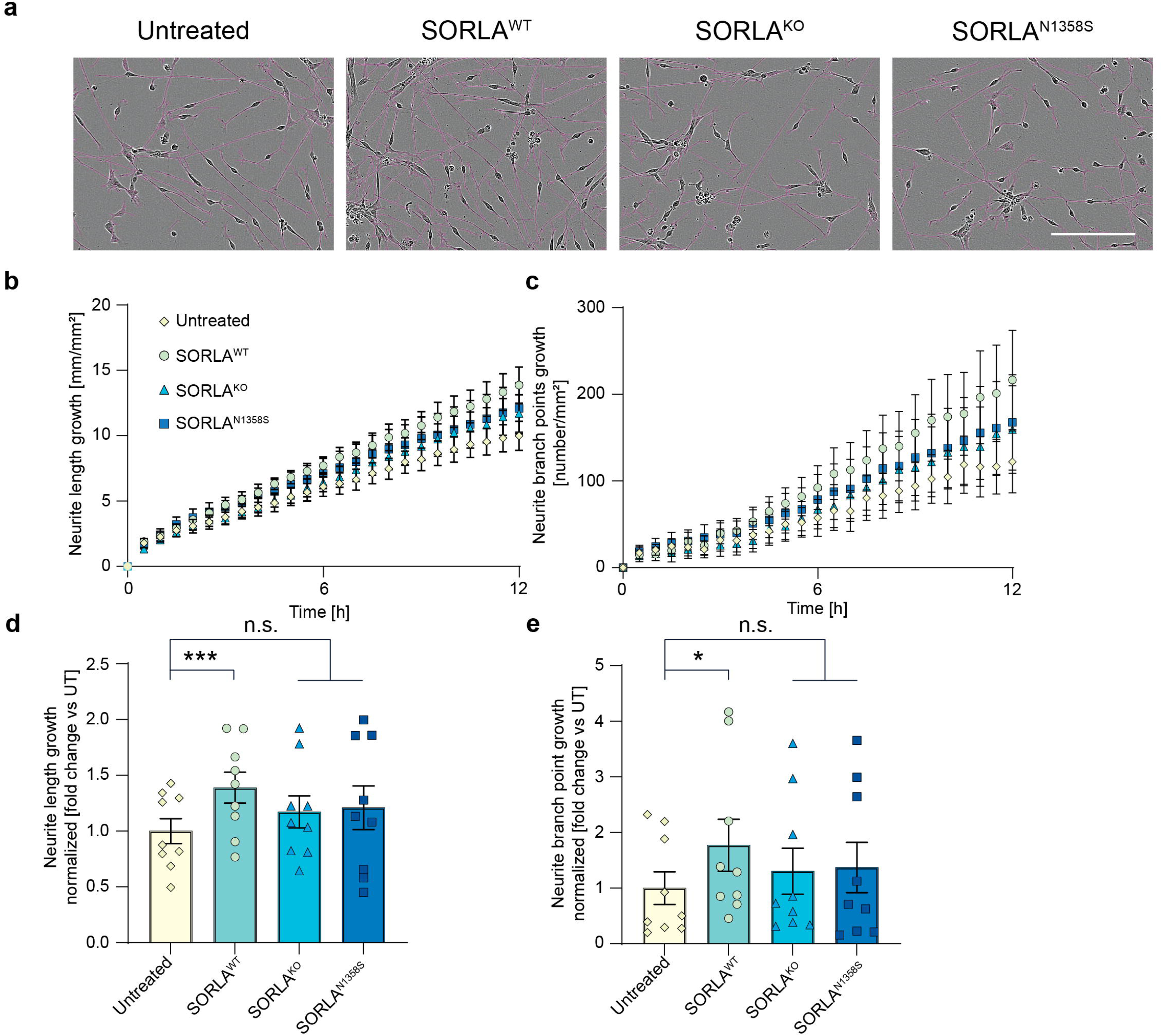
Loss of SORLA or expression of SORLA^N1358S^ reduces the neurotrophic potential of exosomes released by human microglia. **(a)** Brightfield images of immature neurons untreated or treated for 12 h with 10^5^ exosomes/well purified from SORLA^WT^, SORLA^KO^, or SORLA^N1358S^ iMG. Neurite projections, identified by the IncuCyte live cell analysis system (neurotrack software), are superimposed on the images as magenta-colored lines. Scale bar: 200 µm. **(b, c)** Time course analysis of neurite growth (b) and branch point numbers (c) in neurons untreated or treated for 12 h with 10^5^ exosomes/well from iMG of the indicated SORLA genotypes. (**d**) Analysis of neurite growth in immature neurons of after 12 h of treatment (as in b; ns, not significant). Data are given as values normalized the untreated control (mean value set to 1.0). (**e**) Normalized comparison of neurite branch point numbers from neurons after 12 h of treatment with exosomes (as in c). Data in d and e represent 3 independent experiments from individual differentiations, with 3 technical replicates each. Statistical significance was tested using one-way ANOVA with Tukey’s correction for multiple testing (***, p < 0.001).

So far, our studies uncovered an important role for SORLA in defining quantity and neurotrophic quality of exosomes released by iMG. Comparable defects of reduced amounts (Fig. 4e-g) and impaired neurotrophic activity (Fig. 5) were seen in iMG expressing SORLA^N1358S^ or lacking the receptor, documenting that this AD-associated receptor variant is a functional null with respect to control of exosomal biogenesis. This conclusion was corroborated by proteomic analyses of exosome content. All in all, 598 proteins were identified by mass spectrometry in exosome preparations from iMG of the three *SORL1* genotypes. Of these, 58 hits were present in the top 100 list of most expressed exosomal proteins (http://microvesicles.org/extracellular_vesicle_markers), documenting faithful representation of the exosomal proteome in our samples (Supp. table 2). Principal component analysis (PCA) confirmed close similarity of the protein content in exosomes from SORLA^KO^ and SORLA^N1358S^ iMG, but a significant difference of both genotypes to the proteome of SORLA^WT^ exosomes (Fig. 6a). Intriguingly, pathway enrichment analysis identified biological terms related to the function of RNA binding proteins as major distinctions in the exosomal protein contents of SORLA^KO^ and SORLA^N1358S^ as compared to SORLA^WT^ derived exosomes. Exemplary terms included RNA binding and translator regulator activity (for molecular pathways, Fig. 6b), cytosolic small ribosomal subunit and ribosome (for cellular components, Fig. 6c), as well as non-coding RNA processing and cytoplasmic translation (for biological processes, Fig. 6d).

**Figure 6:**
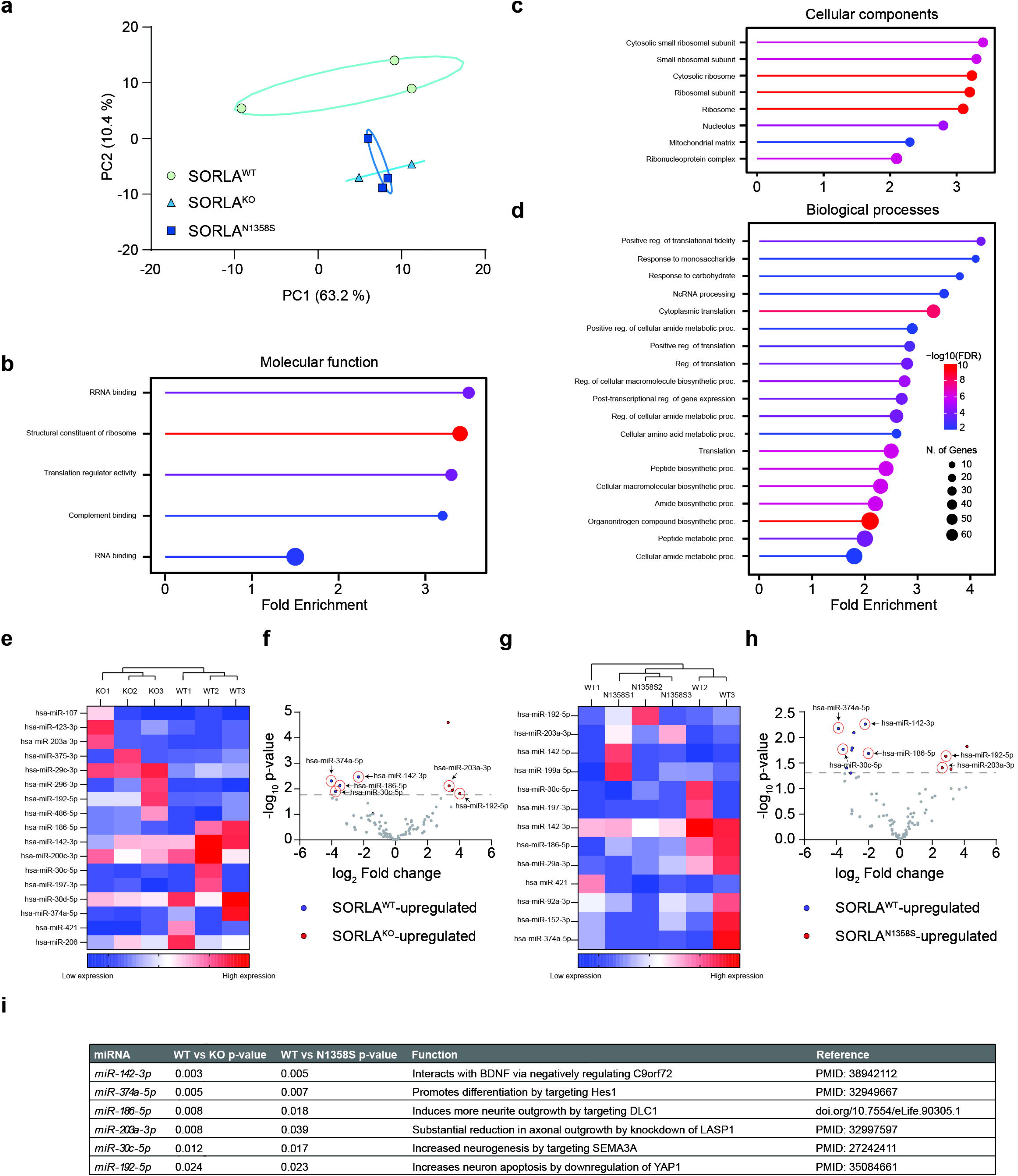
Loss of SORLA or expression of SORLA^N1358S^ impairs inclusion of microRNAs and RNA binding proteins in microglia-derived exosomes. (**a**) Principal component analysis of exosomal proteomes documents comparable protein composition in exosomes secreted from SORLA^N1358S^ and SORLA^KO^ as compared to exosomes from SORLA^WT^ iMG. Data are from 2-3 independent exosome preparations per genotype group. (**d-d**) Gene Ontology Enrichment procedure identifying the most significantly upregulated pathways in the proteome dataset of SORLA^WT^ as compared to SORLA^N1358S^ and SORLA^KO^ derived exosomes. Shown are the most upregulated pathways related to the biological terms molecular pathways (b), cellular components (c), and biological processes (d). (**e-h**) Differential expression analyses of miRNA identified in exosomes from SORLA^N1358S^, SORLA^KO^, and SORLA^WT^ iMG. Analyses were performed using a negative binomial generalized linear model for count-based sequencing data ^55^. miRNAs with p value < 0.05 were considered as significantly changed between genotypes. Heatmaps and volcano plots for comparisons of SORLA^WT^ vs. SORLA^KO^ (e, f), and SORLA^WT^ vs. SORLA^N1358S^ (g, h) are given. miRNAs with p values < 0.05 were determined as significantly changed. The top most differentially expressed miRNA identified in all three genotypes are highlighted by red circles. (**i**) List of top differentially expressed miRNAs in SORLA^WT^ as compared to SORLA^KO^ or SORLA^N1358^ microglia-secreted exosomes. Established functions in control of neuronal outgrowth and maturation are stated.

To interrogate the relevance of altered RNA binding protein composition for changes in neurotrophic potential, we performed bulk RNA sequencing in exosomal preparations from SORLA^WT^, SORLA^KO^, and SORLA^N1358S^ iMG. We focused our analysis on microRNAs that have gained attention as components of exosomes, modulating cell-to-cell communication and driving proliferation and maturation of recipient cells ^37,38^. Using microRNA sequencing strategies, we identified a total of 117 unique microRNAs in exosomes of all three genotypes. Seventeen microRNAs were differently expressed (p < 0.05) when comparing SORLA^WT^ with SORLA^KO^ exosomes (Fig. 6e and f) and 13 when comparing SORLA^WT^ with SORLA^N1358S^ exosomes (Fig. 6g and h). Heatmaps of differential expression data validated the homogeneity of the exosome preparations by grouping SORLA^WT^, SORLAKO, and SORLA^N1358S^ genotypes, individually (Fig. 6e and g). Among the microRNAs up- or downregulated in both SORLA^KO^ and SORLA^N1358S^ when compared to SORLA^WT^ individually, 6 microRNA species were identified as similarly dysregulated in both SORLA^KO^ and SORLA^N1358S^ exosomes (Fig. 6f and h). Of those, all microRNAs increased in levels in SORLA^WT^ exosomes have been ascribed pro-neurotrophic actions, while those with increased levels in SORLA^KO^ and SORLA^N1358S^ exosomes had been associated with functions in limiting neuronal growth and maturation (Fig 6i).

## DISCUSSION

Our studies identified a novel role for the intracellular sorting receptor SORLA in the formation of functional exosomes by neuronal and microglia cell types. This function is lost in an inheritable mutation N1358S, resulting in decreased amounts and impaired neurotrophic potential of exosomes released by cells.

Numerous coding *SORL1* variants have been identified in patients with AD ^7,39^. While the relevance of *SORL1* as a novel disease gene in AD is undisputed, hundreds of coding gene variants in individuals with absent or uninformative short pedigrees makes it difficult to tell disease-causing from common receptor variants ^40^. *In silico* analyses have been used to predict coding variants likely damaging to the receptor structure (www.biorxiv.org/content/10.1101/2023.02.27.524103v1). Not surprisingly, these analyses favor identification of mutations that have a profound effect on the three-dimensional architecture of the receptor polypeptide. Consequently, many mutations studied experimentally globally affect receptor folding, maturation, or trafficking ^10–13^. While these findings indicate a faulty receptor structure as a major cause of AD pathology, they provide limited information on molecular receptor actions essential for brain health. Still, one established receptor activity, clearly substantiated by functional analyses of coding variants, concerns its role in sorting of APP between endosomal compartments and the neuronal cell surface ^10–13^. This sorting path acts neuroprotective as it prevents breakdown of the precursor protein into noxious Aβ peptides in endosomes.

Rather than focusing on genetic loss of known receptor actions, we used an unbiased screening approach to interrogate alterations in a mutant receptor interactome. Using a strategy not constrained by prior hypotheses, we aimed to uncover novel causes of receptor (mal)function relevant to AD. To that end, we focused on mutation N1358S as it maps to the cluster of CR, a well-known ligand binding domain originally found in the low-density lipoprotein receptor and related endocytic receptors ^41^. Also, our *in silico* analysis predicted little impact of this mutation on receptor folding, that would indiscriminately affect all receptor actions (Fig. 1b). Surprisingly, our unbiased screen suggested alterations in the receptor interactome related to biogenesis of extracellular vesicles as the main biological process impacted by N1358S (Fig. 2f). Functional studies in neuroblastoma cells (Fig. 2g-h) as well as in iPSC-derived human neurons (Fig. 3) and microglia (Figs. 4 and 5) fully corroborated a so far unknown role for SORLA in exosome formation and neurotrophic action. Our interactome studies do not distinguish direct from indirect SORLA targets, but the documented profound proteome changes provide an explanatory model for the diminished ability of SORLA^N1358S^ to support cellular production of exosomes.

The biogenesis of exosomes is a complex process tightly connected to vesicular trafficking in endocytic compartments ^30,31^. At present our studies do not reveal a distinct molecular mechanism whereby SORLA guides exosome formation and content. However, its ability to sort proteins between endo-lysosomal compartments and the cell surface ^42^ strongly argues for receptor-dependent sorting of cargo as an important process in defining quantity but also quality of exosomes released by cells. This assumption is supported by colocalization of SORLA with exosomal markers in microglia (Suppl. Fig. 2). Also, this model is supported by prior work from others, identifying structural and functional alterations in endo-lysosomal organelles in neurons and microglia as consequences of SORLA deficiency ^19–21,27^. Exosomal defects in cells expressing mutant SORLA mirror those seen in cells lacking the receptor. Shared phenotypes include a reduction in the amount and neurotrophic quality of exosomes (Figs. 3, 4, and 5) as well as changes in protein and microRNA content (Fig. 6) when compared to exosomes released by wildtype cells. Thus, SORLA^N1358S^ appears as a functional null with respect to this unique receptor activity.

Exosomes play crucial roles in trafficking of proteins, non-coding RNA, and metabolites in the extracellular space, maintaining long distance communication between cells ^43^. In our studies, defects in exosome release are seen in both neurons and microglia lacking SORLA or expressing the N1358S variant, documenting a generalized role for the receptor in formation of exosomes in multiple cell types. Still, this function may be particularly relevant in microglia, where SORLA expression is hypothesized to bare particular importance for AD pathology ^19,25^. In line with this hypothesis, exosomes released by SORLA^N1358S^-expressing microglia fail to promote neurite growth and branching, indicative of a loss of neurotrophic capabilities. microRNA are important determinators of the neurotrophic action of exosomes ^44–46^. Thus, a diminished content of microRNAs and RNA binding proteins in exosomes from mutant SORLA microglia may well be held responsible for their lack of neurotrophic qualities (Fig. 6).

Like other coding mutations identified in *SORL1* thus far, N1358S is a rare variant. Yet, defects in microglia-neuron cross talk through exosomes seen with this mutant may well bare broader significance for a role of the receptor in control of aging brain health and AD. Thus, a number of additional variants have been identified in the CR cluster, which may also have the potential to impact interaction of the receptor with the exosome biogenesis machinery ^39,47–49^. More importantly, this receptor function will be lost in the many *SORL1* variants that abrogate receptor folding and maturation.

## METHODS

### Generation and characterization of SORLA^N1358S^ expressing SH-SY5Y cell lines

The N1358*S* mutation was introduced by site-directed mutagenesis into the human *SORL1* cDNA inserted in expression vector pcDNA3.1zeo+ as detailed in supplementary methods. Functional analysis by surface plasmon resonance analysis ^3^ and EVILFIT ^50,51^ followed published protocols with modifications described in supplements.

### SILAC-based interactome studies

SH-SY5Y lines overexpressing SORLA^WT^ or SORLA^N1358S^ were metabolically labeled for three weeks by culture in SILAC labeling base medium supplemented with normal (“light”) L-arginine (Arg0; Sigma-Aldrich, A6969) and L-lysine (Lys0; Sigma-Aldrich, L8662) or with “heavy” isotope variants Arg10 (^13^C_6_,^15^N_4_; Sigma-Aldrich, 608033) and Lys8 (^13^C_6_,^15^N_2_; Silantes, 211604102). Labeled cell lines were lysed in buffer (Tris-HCl, pH 8.0, 20 mM NaCl, 0.6% w/v sodium deoxycholate, 0.6% w/v NP-40) containing protease inhibitor (Roche, 11697498001). Home-made goat anti-human SORLA IgG or non-immune IgG were coupled to Pierce™ NHS-Activated Magnetic Beads (Thermo Fisher, 88826) and used for immunoprecipitation. After elution with 6 M guanidium-HCl for 10 min at 70 °C, eluted proteins were precipitated using ethanol and digested with trypsin. Peptide extracts were purified and stored on stage tips according^52^. Mass spectrometer-based sequencing of the peptides and bioinformatic analysis of data are described in the supplements.

### Generation and differentiation of iPSC lines

Human-induced pluripotent stem cell (iPSC) line hpscreg.eu/cell-line/BIHi043-A was used as the wildtype control. Isogenic *SORL1*-deficient line hpscreg.eu/cell-line/BIHi268-A-18 and isogenic clone hpscreg.eu/cell-line/BIHi268-A-44, homozygous for *SORL1^N1358S^*, were generated from the parental line by CRISPR/Cas9-mediated genome editing ^53^, and differentiated into cortical neurons ^24^ or microglia ^28^ using established protocols. Details of genome editing, karyotype validation, as well as differentiation protocols are given in the supplementary methods.

### Purification and functional characterization of exosomes

Exosomes were purified from supernatants of cells using published protocols ^22^ as detailed in the supplements. Exosomes were subjected to nanoparticle tracking analysis (NTA) using a NanoSight NS300 system (Malvern Panalytical). For measurement, exosome samples were diluted 1:100 in PBS, mixed, and injected into the sample chamber. The measurement script comprised temperature control at 23°C, followed by a 20 s flush at a flowrate mark of 1000. Next, sample advancement was stabilized over 120 s at flowrate mark 10. Recordings were captured continuously during a steady flow at flowrate mark 10 with five 60 s recordings separated by a 5 s lag time between samples. Videos were collected and analyzed using NanoSight software (version 3.3 and 3.4). Automatic settings were used for the max jump mode, minimum track length, and blur setting. With these settings, the max jump distances were between 19.0 and 22.7. Camera level and detection thresholds were adjusted according to sample composition to ensure optimal sensitivity. The camera level was set to a maximum of 16 to ensure maximum sensitivity for small vesicles. Detection threshold was set to 5. Further analysis of exosome preparation by mass spectrometry-based proteomics and electron microscopy are detailed in supplementary methods.

### Neurite outgrowth and maturation assays

Immature neurons were generated from human iPSC line BIHi005-A-24, engineered to stably carry a doxycycline (dox)-inducible expression construct for transcription factor neurogenin 2 (NGN2) in the *AAVS1* locus ^54^. We wish to acknowledge Dr. M. Peitz (UKB) for providing plasmids and the Stanford University Cardiovascular Institute Biobank, especially Prof. J. Wu, for SCVI 111 fibroblasts used to generate this line.

On day −1, 80% confluent iPSC cultures were disassociated with 0.5 mM EDTA/PBS and seeded on Matrigel-coated 6-well plates at a density of 450,000 cells/well in E8 Flex supplemented with 10 μg/ml Y27632. On day 0, the medium was changed to neuronal induction medium (NIM) (DMEM/F-12; Gibco, 11330-032) and 1 % N2 (Gibco, 17502048) supplemented with 2 µg/ml dox. On day 1, the medium was changes to fresh NIM. On day 2, cells were dissociated with TrypLE (Gibco, 12604013) for 10 min at 37°C. Subsequently, the cells were plated on poly-ornithine/laminin-coated 96 wells at a density of 20,000 cells/well. The medium was changed to Neurobasal medium (Gibco, 21103049) with 2 % B-27 supplement (Gibco, 17504044), 1% GlutaMAX (Gibco, 35050061), and 10 ng/ml BDNF (Gibco, P23560) supplemented with 2 µg/ml dox and 10 μg/ml Y27632. On day 3, the medium was changed to fresh medium supplemented with 2 µg/ml dox and 10 µM -[N-(3,5-Difluorophenacetyl)-L-alanyl]-S-phenylglycine t-butyl ester (DAPT; Merck, D5942). After 4 h, the cells were treated with exosomes for assessment of neurite growth and maturation.

Immature neurons prepared as described above were treated with either untreated or treated with 10^5^ exosomes purified from the supernatants of SORLA^WT^, SORLA^KO^, or SORLA^N1358S^ microglia. Exosomes had been prediluted in PBS in accordance with NTA measurements at a concentration of 2×10^7^ exosomes/ml. Five µl exosome dilutions were added to 195 µl media in each well. Then, the 96-well plate was live-imaged for 12 h using an IncuCyte S3/SX1 G/R live cell imaging and analysis system (Sartorius, Gottingen) with 4 pictures taken for each well at 20x magnification every 30 min. Data analysis was performed using the neurotrack software package (Sartorius, Gottingen) for three wells for each condition. Features analyzed included neurite outgrowth (mm/mm^2^) and neurite branch points (number/mm^2^). For each well, an average of 4 pictures was analyzed for each time point. Data were normalized to show the increase in neurite length and number of branch points from time point t = 0 h. Comparison at 12 h between untreated and exosome-treated conditions were normalized to the untreated condition as baseline.

### Statistical analyses

The number of n represents biological replicates collected from a minimum of 2 independent differentiation experiments. For co-localizations, n is the number of cells analyzed from multiple independent experiments. Statistical analyses were conducted in GraphPad Prism version 10. Data are presented as mean ± SEM with details of statistical analyses specified in figure legends.

## Contributions

KJ-M, I-MR, KM, OMA, MS, and TEW conceptualized the study. KJ-M, I-MR, JPG, KM, PLO, MG-W, MK, NST, MTS, DSS, JP, SD, OMA, MS, and TEW developed the methodology. KJ-M, I-MR, JPG, KM, PLO, MG-W, MK, NST, MTS and JP made the investigations. KJ-M, JPG, OMA, and TEW designed the visualization of data. KJ-M, and TEW secured funding acquisition. KJ-M and TEW undertook project administration. KJ-M and TEW performed project-related supervision and wrote the original draft of the manuscript. KJ-M, I-MR, JPG, KM, PLO, MG-W, MK, NST, MTS, DSS, JP, SD, OMA, MS, and TEW reviewed and edited the manuscript.

## Acknowledgements

We are indebted to T. Pasternack, K. Kampf, A. Høiland, and K.M. Pedersen for expert technical assistance. Studies were funded by the AFI (#18003) and Novo Nordisk Foundation (NNF18OC0033928) to TEW, and by the Lundbeck Foundation (R380-2021-1326) to KJM.

## Data availability

The mass spectrometry proteomics data have been deposited to the ProteomeXchange Consortium via the PRIDE partner repository with the dataset identifier PXD061828. All other data will be made available upon reasonable request to the corresponding authors.

## List of Abbreviations

(SORLA): sortilin-related receptor with A-type repeats
(APP): amyloid precursor protein
(Aβ): amyloid-β peptides
(AD): Alzheimer’s disease
(SNPs): single nucleotide polymorphisms
(CR): complement-type repeats
(SPR): surface plasmon resonance
(WT): wildtype
(RU): response units
(SILAC): stable isotope labeling by amino acids in cell culture
(IP): immunoprecipitated
(NTA): nanoparticle tracking analysis
(iMG): induced human microglia

