## Supplements for "Familial Alzheimer disease mutation identifies novel role of SORLA in release of neurotrophic exosomes"

**SUPPLEMENTARY INFORMATION LIST OF CONTENT**

1) Supplementary methods

2) Supplementary references

3) Supplementary tables 1 and 2

4) Supplementary figures 1 and 2, and legends

### 1) SUPPLEMENTARY METHODS

#### Alphafold analysis

The structural implications of mutation N1358S were evaluated using alphafold 2 (1). In brief, the wildtype and mutant sequences of human SORLA (amino acids K497 to R1883) were loaded into neurosnap (2) and visualized using PyMOL and a ribbon representation of the predicted structure with the mutated site highlighted using spheres.

#### Generation SORLA$^{N1358S}$ expressing SH-SY5Y cell lines

The N1358S mutation was introduced by site-directed mutagenesis into the human *SORL1* cDNA inserted in expression vector pcDNA3.1zeo+. To do so, the *SORL1* cDNA was amplified in two fragments using primers A (5′-AAGAGAATGTCCACAGCTGG-3′) and B (5′-GTTTTCGCAG**C**TGGCTTCATCAGAATAATC-3′), as well as primers C (5′-ATTATTCTGATGAAGCCAG**C**TGCGAAAACC-3′) and D (5′-CATCCTGGCAATCTTGGTGG-3′), with primers B and C carrying the mutation to be introduced (given in bold). Subsequently, PCR products AB and CD where re-amplified using primers A and D. The mutated His-tagged SORLA ectodomain used for SPR analysis was generated using the same cloning strategy. Plasmid transfections into SH-SY5Y cells were performed using Lipofectamine™ 2000 (Thermo Fisher) according to manufacturer's instructions and stable SORLA overexpressing lines selected using 50 µg/ml zeocin.

**Surface plasmon resonance (SPR) analysis**

SPR analysis was carried out using a BIAcore2000 system as described before (3). In short, pCepPu expression constructs encoding the entire hexa-His-tagged ectodomains of $SORLA^{WT}$ or $SORLA^{N1358S}$ were transfected into HEK293-EBNA cells and proteins purified from conditioned media 48 hours later using standard $Ni^{2+}$ affinity chromatography. Purified ectodomains of SORLA were immobilized on a CM5 Biacore chip at a density of 56 $fmol/mm^2$. The sensor chip was incubated with 100 nM purified hexa-His-tagged ectodomain of $APP^{695}$ in running buffer (10 mM Hepes, pH 7.4, 150 mM NaCl, 5 mM $CaCl_2$, 0.005% Tween 20). The difference in response signals between SORLA-coated and non-coated control chips was recorded as relative response units (RU).

**EVILFIT**

To derive binding kinetics from SPR data considering any possible multivalent interaction, we followed established procedures (4). In brief, according to Kitov's and Bundle's model (5), the multivalent binding event can be mathematically modeled as a set of parallel reactions. The SPR experiment permits these reactions to be approximated as an ensemble of multiple pseudo-first-order reactions. The minimal set of such a 1:1 reaction ensemble can be calculated using the EVILFIT algorithm (6). Two-dimensional fits of the SPR data were generated on the MATLAB 2012a platform (Mathworks) using fitting tool EVILFIT version 3 (7, 8). Input values matched the start and end injection time and a concentration of 100 nM $APP^{695}$. The association phase was fitted from t = 'injection start' plus 1 s to t = 'injection end' minus 5 s. The dissociation phase was fitted from t = 'injection end' plus 5 s to t = 'injection end' plus 500 s. The operator-set boundaries for the distributions were uniformly set to limit $K_D$ values in the interval from $10^{-11}$ to $10^{-2}$ M, and $k_{off}$ values in the interval from $10^{-7}$ to $10^{-1}$ $s^{-1}$ to ensure comparable and best quality fits reflected in a high signal to the root mean square distance (rmsd) ratio. The distribution P (ka, KA) was calculated using the discretization of equation:

$$R_{total} = \int_{K_{A_{min}}}^{K_{A_{max}}} \int_{k_{a_{min}}}^{k_{a_{max}}} R\left(k_a, K_A, C_{analyte}\right) P(k_a, K_A) dk_a dK_A$$

in a logarithmic grid of (ka,i, KA, i) values with 15 grid points distributed on each axis. This was done through a global fit to association and dissociation traces at the given analyte concentration. Tikhonov regularization was used at a confidence level of P = .95 to determine the most parsimonious distribution that is consistent with the data, showing only features that are essential to fit the data (9). The resulting 2D-distribution of minimally required 1:1 interaction to model the data is plotted in a grid of resulting $K_D$ and $k_{off}$ rates. The total rmsd between total fit and experimental data was 0.4 and 0.6 % for $SORLA^{WT}$ and $SORLA^{N1358S}$, respectively (Fig. 1g-e).

**SILAC-based interactome studies**

SH-SY5Y cell lines stably overexpressing $SORLA^{WT}$ or $SORLA^{N1358S}$ were metabolically labeled with heavy or light isotope amino acids for three weeks by culture in SILAC labeling base medium supplemented with normal ("light") L-arginine (Arg0; Sigma-Aldrich, A6969) and L-lysine (Lys0; Sigma-Aldrich, L8662) or with "heavy" isotope variants Arg10 ($^{13}C_6$, $^{15}N_4$; Sigma-Aldrich, 608033) and Lys8 ($^{13}C_6$, $^{15}N_2$; Silantes, 211604102). Subsequently, labeled cell lines were lysed in buffer (Tris-HCl, pH 8.0, 20 mM NaCl, 0.6% w/v sodium deoxycholate,

0.6% w/v NP-40) containing protease inhibitor (Roche, 11697498001). Home-made goat anti-human SORLA IgG or non-immune IgG were coupled to Pierce™ NHS-Activated Magnetic Beads (Thermo Fisher, 88826) according to manufacturer's instructions and used for immunoprecipitation. After elution with 6 M guanidium-HCl for 10 min at 70 °C, eluted proteins were precipitated using ethanol and digested with trypsin. The digestion was stopped by acidifying each sample to pH < 2.5 by adding 10% trifluoroacetic acid solution. The peptide extracts were purified and stored on stage tips according to (10).

Peptides were eluted using Buffer B (80% Acetonitrile and 0.1% formic acid) and organic solvent was evaporated using a speedvac (Eppendorf). Samples were diluted in Buffer A (5% acetonitrile and 0.1% formic acid). Peptides were separated on a reversed-phase column with 136 min gradient with a 250 nl/min flow rate of increasing Buffer B concentration on a High Performance Liquid Chromatography (HPLC) system (Thermo Fisher Scientific). Peptides were ionized using an electrospray ionization (ESI) source (Thermo Fisher Scientific) and analyzed on a Q-exactive plus Orbitrap instrument (Thermo Fisher Scientific). The mass spectrometer was run in data dependent mode selecting the top 10 most intense ions in the MS full scans, selecting ions from 300 to 1700 m/z (Orbitrap resolution: 70,000; target value: 1,000,000 ions; maximum injection time of 120 ms). The resulting MS/MS spectra from the Orbitrap had a resolution of 17,500 after a maximum ion collection time of 60 ms with a target of reaching 100,000 ions.

The resulting raw files were analyzed using MaxQuant software version 1.5.2.8 (11). Default settings were kept except that 'match between runs' and 're-quantify' was turned on. Lys8 and Arg10 were set as labels and oxidation of methionines and N-terminal acetylation were defined as variable modifications. Carbamidomethyl of cysteines was set as fixed modification. The *in silico* digests of the human Uniprot database (2015-12) and a database containing common contaminants were done with Trypsin/P. The false discovery rate was set to 1% at both the peptide and protein level and was assessed by in parallel searching a database containing the reversed sequences from the Uniprot database. The mass spectrometry proteomics data have been deposited to the ProteomeXchange Consortium via the PRIDE partner repository (12) with the dataset identifier PXD061828.

**Over-representation analysis (ORA) of SILAC data sets**

ORA of the SILAC data sets was performed using Genekitr. Gene names of proteins with decreased interaction with SORLA$^{N1358S}$ as compared to SORLA$^{WT}$, comprising a total of 152 proteins from SH-SY5Y clones 59 and 76, were used as input. Gene hits were compared to a gene list from *Homo sapiens* comprising 20,039 genes. Gene ontology (GO) analysis was performed using the cellular component gene set and a p-value and adjusted p-value cut-off of 0.05. Q-value cut-off was likewise set to 0.05 and Benjamini-Hochberg (BH) test was applied to calculate adjusted p value of GO terms. The minimal gene set was set to 10 and the maximum gene set to 5000.

**Generation of *SORL1*-deficient and N1358S mutant iPSC lines**

Human induced pluripotent stem cell (iPSC) line BIHi043-A was used as the wildtype control line in this study. The isogenic *SORL1*-deficient line BIHi268-A-18 was generated from the parental line by CRISPR/Cas9-mediated genome editing, as described (13) using a gRNA targeting exon 1 in *SORL1* (5´-CAGTAGCGTTCGCCCGAACA-3´). An 8 bp frameshift deletion predicted to cause inactivation of *SORL1* in a targeted cell clone was confirmed by Sanger sequencing using primers 5´-AGAAAGTGCGCGAAAGGGA-3´ (forward) and 5´-

AAAACTGCTCACCTGTCCGT-3′ (reverse). Likewise, the parental BIHi043-A line was used to generate an isogenic iPSC clone BIHi268-A-44, homozygous for the *SORL1*$^{N1358S}$ mutation. In detail, an optimized single guide RNA (sgRNA) targeting *SORL1* at the desired mutation site was designed and synthesized by Integrated DNA Technologies (https://eu.idtdna.com). The sgRNA sequence was 5′-TATTCTGATGAAGCCAACTG-3′. A single-stranded oligodeoxynucleotide (ssODN) donor template was designed to change codon AAC to AGC and synthesized as an Ultramar DNA oligo by IDT, with the sequence 5′-GCATCACTGTTAACTAGAGGATTAGGATGTGAGGAGGCAGACTGGACATCTTACCGCAGCTTGCTTCATCAGAATAATCGCCGCAATCATCCATCCCGTCGCACTTCCAAATCAAACTGA-3′. In addition, the sgRNA sequence introduced a silent mutation in the preceding codon for alanine from GCC to GCA. This ‘CORRECT’ method is commonly used to improve homology-directed repair accuracy by preventing re-cutting of the targeted site by Cas9, facilitating precise and scarless genome editing (14). Ribonucleoprotein (RNP) complexes were assembled by mixing 1.5 µg of Cas9 protein with 360 ng of gRNA and incubating for 10 minutes at room temperature. Next, RNP complexes, along with the ssODN template, were added to a suspension of $1x10^5$ cells using the Neon transfection System (Thermo Fisher). Following electroporation, the cells were plated in a 6-well plate using StemFlex medium supplemented with CloneR (Stemcell Technologies). Three days post-transfection, the cell population was tested for the presence of the desired mutation by PCR amplification and Sanger sequencing. Primers used were 5′-ACCACTGCTGCTTCGAC-3′ and 5′-CCCTTCAAGACTCAACTCGTAT-3′. Subsequently, single-cell cloning of the edited cells was carried out. Single-cell clones were verified by Sanger sequencing and correctly targeted clones expanded and quality-controlled by SNP karyotyping as described (15). All cell lines were routinely tested negative for mycoplasma.

**Differentiation of iPSC into human neurons and microglia**

For maintenance, iPSC lines were cultured on Matrigel (Gibco, 356324) coated 6-well plates in Essential 8™ Flex Medium (Gibco, A2858501). The culture medium was changed every second day and cells were passaged in clusters every 3-4 day at a density of 80% using 0.5 mM EDTA/PBS. The iPSC lines were maintained for 3-10 passages before starting a new differentiation experiment.

Induced neurons (iNs) were generated from iPSCs as described previously (16). Briefly, iPSCs were dissociated with accutase and replated at a density of 5.5 x $10^4$ cells/$cm^2$ in Matrigel-coated 24- or 12-well plates or 4.3 x $10^4$ cells/$cm^2$ in Matrigel-coated 6-well plates and 100 mm-dishes using E8 Flex media supplemented with 10 µM Rock inhibitor (Y27632; Cayman Chemical). Next day (day -1), iPSCs were transduced with lentivirus vectors in fresh E8 Flex medium containing 7 µg/ml Polybrene (Millipore). One day after infection (day 0), the medium was replaced with F12-N2 (DMEM/F12, 1% N2, 1% NEAA (Gibco)), containing 2 µg/ml doxycycline, 10 ng/ml human BDNF (R&D Systems), 10 ng/ml human NT3, and 0.2 µg/ml mouse laminin (Thermo Fisher). Doxycycline was retained in the medium through the experiment. On day 1, the medium was replaced with fresh F12-N2 medium containing 0.8 µg/ml puromycin to select vector expressing *Ngn2* (kindly provided by T. Sudhof, Standford Medicine). At day 2, half of the F12-N2 medium was replaced with NB-B27 medium. From day 3 on, half of the medium was replaced by NB-B27 medium every second days. Induced neurons were kept in culture for 11 days prior to exosome collection.

Differentiation of iPSC into induced microglia (iMG) was done according to published protocols (17). In brief, iPSCs were differentiated into hematopoietic progenitors (HP) using the STEMdiff™ Hematopoietic Kit (Stem Cell Technologies, #05310). To do so, iPSCs at a confluency of 70-80% were passaged with ReLeSR™ (Stem Cell Technologies, 05872) in E8 flex containing Matrigel coated 6-well plates (day -1). Cell clusters of 100 cells were seeded at a density of 50-100 clusters per well. At day 0, in wells with a total of 40-80 clusters, E8 flex medium was replaced with 2 ml of medium A. On day 2, 1 ml of medium A was added to the well. On day 3, the medium was replaced with 2 ml of medium B. At days 5, 7, and 10, 1 ml medium B was added. At day 12, the media was gently resuspended in the wells using a 5 ml serological pipette to increase the yield of HPs. Collected HPs were centrifuged at 300x g for 5 min, resuspended in microglia differentiation medium (DMEM/F-12, Gibco) containing 2x Insulin-Transferrin-Selenite (Thermo Fisher, 41400045), 2x B27 (Thermo Fisher, 17504001), 0.5x N2 (Thermo Fisher, 17502048), 1x GlutaMAX (Thermo Fisher, 35050038), 1x NEAA (Thermo Fisher, 11140035), 400 µM monothioglycerol (Sigma, M1753), 5 µg/ml insulin (PromoCell, C-52310), 100 ng/ml IL-34 (PeproTech, 200-34), 50 ng/ml TGFβ1 (PeproTech, 100-21C), and 25 ng/ml M-CSF (PeproTech, 300-25). Cells were seeded at a density of 200,000 cells each in Matrigel coated 6-well plates. On days 14, 16, 18, 20, and 22, 1 ml of microglia differentiation medium was added to the wells. At day 24, 5 ml of medium was transferred to a 15 ml tube to spin down the cells at 300x g for 5 min. The supernatant was removed and the cell pellet resuspended in 1 ml differentiation medium and returned to the well. This procedure was repeated daily for days 26 through 33. At day 35, the cell pellets were resuspended in microglia maturation medium, containing 100 ng/ml CD200 (Elabscience, E-PKSH032840) and 100 ng/ml

CX3CL1 (PeproTech, 300-31), for further maturation. On day 37, 1 ml of microglia maturation medium was added to the well. Between days 38-42, iMG were used for functional studies.

**Expression analyses**

Transcript levels in iPSC, iNs and iMG were analyzed by quantitative RT-PCR. In brief, total RNA was isolated using the RNeasy Plus Micro kit (Qiagen) and reverse transcribed to cDNA with high-capacity RNA to cDNA kit (Applied Biosystems). The cDNA samples were subjected to quantitative (q) RT-PCR using TaqMan Gene Expression Assays *SOX2* (Hs01053049_s1), *OCT4* (Hs00999632_g1), *NANOG* (Hs02387400_g1), *SORL1* (Hs00983770), *IBA1* (Hs00610419), *P2RY12* (Hs01881698), *CX3CR1* (Hs01922583_s1), *TREM2* (Hs00219132_m1), *HPRT1* (Hs02800695_m1), *GAPDH* (Hs02758991_g1), *TBP* (Hs00427620_m1), *MAP2* (Hs00258900_m1). Relative gene expression was determined with the cycle threshold (CT) comparative method ($2^{-ddCT}$). Data were normalized to reference genes *GAPDH*, and *HPRT1,* and fold changes relative to levels in the given controls were determined as stated in the respective figure legends.

Protein levels were evaluated by standard Western blot analyses using primary and horseradish peroxidase-conjugated secondary antibodies (Sigma-Aldrich) and the SuperSignal West Femto Maximum Sensitivity Substrate (Thermo Fisher). Primary antibodies were mouse anti-SORLA (BD611861, BD Biosciences) and mouse anti-GAPDH (gtx627408, GeneTex).

**Immunocytochemistry**

Induced microglia (iMG) were seeded onto non-coated 12 mm glass coverslips in a 24-well plate and allowed to attach overnight. The next day, cells were washed in PBS once and fixed with 4% PFA/PBS for 10 min at room temperature. For all immunostainings, cover slips were washed in

PBS, incubated with blocking buffer (5% normal donkey serum, 0.1% Triton X-100 in PBS) at room temperature for 30 min, followed by incubation with primary antibodies in blocking buffer at 4°C overnight. The next day, cover slips were washed in PBS, incubated with secondary antibodies in blocking buffer (1:500) for 2 hours at room temperature, counterstained with DAPI (1:3000, 10236276001, Roche), and mounted in fluorescent mounting media (DAKO). Primary antibodies were in-house goat and rabbit anti- human SORLA antisera, as well as commercially available antibodies directed against IBA1 (ab5076, Abcam), P2RY12 (HPA014518, Sigma), TREM2 (MABN755, Merck), CD81 (TAPA 5A6, Biolegend), CD9 (H19a, Biolegend), and CD63 (H5C6, BD Pharmingen). Co-localization of immunostained SORLA and exosomal biogenesis markers were analyzed by Pearson's correlation coefficient using FIJI (ImageJ) and the Bioimaging and Optics Platform version of JaCoP (18).

**Purification of exosomes**

Exosomes were purified from supernatants of cells using published protocols (19). In brief, 80% confluent cultures of SH-SY5Y cells grown in T75 culture flasks were washed twice in PBS before 20 ml of serum-free DMEM was added and conditioned for 16 h. For collection of exosomes from iN, media was removed on day 11 of differentiation and 8 ml NB-B27 media was added to be conditioned for 16 h. For collection of exosomes from iMG, cells were harvested on day 37 and washed twice in PBS before 500,000 cells were seeded into a 150 mm non-coated dish (734-2322, VWR) with 20 ml differentiation media to be conditioned for 16 h. All purification steps were performed on ice to assure integrity of exosomes. Cell supernatants were spun twice at 1,000g for 5 min, discarding the pellet to remove cell debris. Next, supernatants were carefully filtered through a 0.22 μm sterile filters (SEM1M179M6, Millipore), rinsed twice with 5 ml ice-cold PBS to remove microvesicles above 200 nm in size. Then, exosomes suspensions were centrifuged at 100,000g for 4 h at 4°C using a pre-cooled Type 45 Ti fixed-angle rotor. Supernatants were discarded and pellets resuspended in 30 ml PBS before re-centrifugation at 100,000g for 2 h at 4°C. Finally, supernatants were removed and exosome pellets dissolved in 100 µl PBS and aliquoted into 10 µl aliquots for storage at -80°C.

**Exosome proteomics**

Purified exosome pellets were lysed in RIPA lysis buffer (25 mM Tris-HCl pH 7.6, 150 mM NaCl, 1% NP-40, 1% sodium deoxycholate and 0.1% SDS) containing complete mini EDTA-free protease inhibitor cocktail and PhosSTOP phosphatase inhibitor cocktail and further sonicated for 15 cycles (30 sec on and 30 sec off). The protein concentration was determined by Pierce Bradford Protein Assay Kit (Thermo Fisher Scientific). For label-free quantification, 20 µg of extracted proteins per sample were fractionated by SDS-polyacrylamide gel electrophoresis on 4–15% Criterion TM TGX™ precast gel. The proteins were subjected to reduction with dithiothreitol and alkylation with iodoacetamide before in-gel trypsin digestion. The resulting peptides were purified with Pierce TM C18 Spin Columns and vacuum dried before MS analysis. Peptide samples were trapped and separated by liquid chromatography (Easy-nLC 1200, Thermo Scientific) using precolumn (Acclaim PepMap 100, 75 µm × 2 cm, Nanoviper, Thermo Scientific) and analytical column (EASY-Spray column, PepMap RSLC C18, 2 µm, 100 Å, 75 µm × 25 cm) in a 110 min gradient of 4–40% acetonitrile in 0.1% formic acid, coupled to the mass spectrometer Q-Exactive HF-X Hybrid Quadrupole Orbitrap (Thermo Scientific, Bremen). Precursor ion mass spectra (MS1) were acquired at 60,000 resolution and the scan range was between 372 to 1800 m/z. MS2 analysis was performed in a data-dependent mode, where the most intense doubly or multiply charged precursors were fragmented. MS2 resolution was set at 15,000. Unassigned and +1 charge state was excluded from fragmentation and a dynamic exclusion of 15 s was used. Proteins were identified and quantified and in at least two out of three replicates using MaxQuant (version 1.5.3.30) against a human sequence database (Homo Sapiens proteome with 20,129 reviewed sequences downloaded from Uniprot database, https://www.uniprot.org/ on December 12th, 2016). Raw intensities extracted by MaxQuant were log(2) transformed, and normalized by median of replicates for label-free quantification. Missing values were replaced by imputation according to normal distribution. Differentially expressed proteins were identified by using unpaired two-tailed Student's t test with the thresholds of ±1.5-fold change over the control and a p-value < 0.05. Gene ontology (GO) enrichment analysis of differentially expressed proteins was performed using ShinyGO 0.82 (https://pmc.ncbi.nlm.nih.gov/articles/PMC7178415/) with default settings. The total list of all proteins identified in exosome samples was used as background.

**Electron microscopy of exosome preparations**

All TEM grids were prepared with a negative staining protocol modified from (20). TEM grids (CF400-Cu) were obtained from Electron Microscopy Sciences (Hatfield, PA) and a PELCO easiGlow™ (Ted Pella) was used for glow discharging. A 5 µl droplet of freshly thawed exosome sample was placed on a freshly peeled piece of Parafilm®. Hereafter, a freshly glow-discharged treated TEM grid (45 s at 25 mA) was placed with the carbon-coated side on the droplet and left to incubate for 10 minutes. Hereafter, two 4 µL droplets of freshly thawed 2% uranyl formate (w/v in DI water) were placed on the Parafilm® piece. Then, the TEM grid was blotted dry using a Whatman® grade 1 filter paper, and immediately after placed on the first uranyl formate droplet, quickly blotted dry, and then left on the second uranyl formate droplet for 20 s. Hereafter, the TEM grid was blotted dry and was ready for imaging.

The TEM grids were imaged using a Tecnai G2 spirit (FEI company) operating at 120 kV, mounted with a TVIPS TemCam F-416, 4k CMOS camera. Illumination was done with e-beam spot size 2, using a $LaB_6$ filament, the condenser aperture in position 3 (100 µm), and objective aperture in position 4 (100 µm). All grids were imaged using 26.000 times magnification to obtain overview images and 150.000 times magnification for closeup images using similar imaging conditions. For every sample, a minimum of 6 images in high and low magnification were obtained at selected locations, with the criterion that the image should contain at least one representative-looking exosome for the sample.

**Bulk RNA sequencing of exosomes**

RNA sequencing was done by EV Genomics (https://www.evgenomics.com/). For RNA library preparation, total RNA was isolated from 100 µl of purified exosome samples using the miRNeasy Serum/Plasma Advanced Kit (Qiagen, Cat. No. 217204) following the manufacturer's protocol. RNA was eluted in 12 µl of RNase-free water and quantified using the Bioanalyzer RNA 6000 Pico Kit (Agilent, Cat. No. 5067-1513). For small RNA library preparations, 5 µl of eluted RNA were processed using the QIAseq miRNA Library Kit (Qiagen, Cat. No. 331505). In brief, 3′ and 5′ adapters were ligated to the RNA, followed by reverse transcription with a primer containing a unique molecular index (UMI). Library amplification was performed using 22 PCR cycles to introduce the sample index. Library quality and quantity were assessed using the Bioanalyzer High Sensitivity DNA Analysis Kit (Agilent, Cat. No. 5067-4626). Finally, libraries were pooled and sequenced on one lane of a NovaSeq Plus 10B flow cell (Illumina, San Diego, CA, USA) using single-end 150 bp sequencing.

Raw sequencing data (FASTQ files) were imported into the CLC Genomics Workbench (QIAGEN RNA-seq Analysis Portal 5.1, https://rnaportal.qiagen.com) for microRNA analysis. Quality control included filtering low-quality reads and removing adapter sequences. Reads were merged using Unique Molecular Indexes (UMIs) to correct for PCR amplification bias. microRNAs were identified, counted, and annotated using miRBase v22 aligned to the Homo sapiens genome (GRCh38.103). Expression levels of mature miRNAs were quantified and normalized using the Trimmed Mean of M-values (TMM) method to adjust for differences in library size. Differential expression analysis was performed using a negative binomial Generalized Linear Model (GLM), appropriate for count-based sequencing data. microRNAs with P-value<0,05 were determined as significantly changed.

### 3) SUPPLEMENTARY TABLES, FIGURES AND LEGENDS

**Supplementary table 1: Proteins identified by over-representation analysis of cellular component pathways in the SORLA interactome**

The table highlights four exemplary cellular component pathways enriched in the receptor interactome identified by comparative SILAC of $SORLA^{WT}$ versus $SORLA^{N1358S}$ (see Fig. 2f). For each component compartment, 10 proteins and their associated protein class are listed. Common proteins identified by SILAC in $SORLA^{N1358S}$ clones 59 and 76 by SILAC are given in bold.

| **Extracellular exosome** | | |
|---|---|---|
| **Gene** | **Protein** | **Protein class** |
| ***YWHAB*** | **Tyrosine 3-monooxygenase** | Scaffold/Adaptor Protein |
| ***RAC1*** | **Ras-related C3 Botulinum toxin substrate 1** | Small GTPase |
| ***RAC2*** | **Ras-related C3 Botulinum toxin substrate 2** | Small GTPase |
| ***MFGE8*** | **Lactadherin** | Oxidoreductase |
| ***HNRNPL*** | **Heterogeneous nuclear ribonucleoprotein L** | RNA processing factor |
| *CLTC* | Clathrin heavy chain | Vesicle coat protein |
| *LDHA* | Lactate dehydrogenase A | Dehydrogenase |
| *RAB5C* | Ras-related protein Rab-5C | Small GTPase |
| *PSMA3* | Proteasome 20S subunit alpha 3 | Protease |
| *CDk1* | Cyclin dependent kinase 1 | Non-receptor Serine/Threonine protein kinase |
| **Extracellular space** | | |
| ***YWHAB*** | **Tyrosine 3-Monooxygenase** | Scaffold/Adaptor Protein |
| ***RAC1*** | **Ras-related C3 Botulinum toxin substrate 1** | Small GTPase |
| ***RAC2*** | **Ras-related C3 Botulinum toxin substrate 2** | Small GTPase |
| ***MFGE8*** | **Lactadherin** | Oxidoreductase |
| ***HNRNPL*** | **Heterogeneous nuclear ribonucleoprotein L** | RNA processing factor |
| ***SEMA3A*** | **Semaphorin-3A** | Membrane-bound signaling molecule |
| *VGF* | Neurosecretory protein VGF | Intercellular signal molecule |
| *CNTN1* | Contactin-1 | Cell adhesion molecule |
| *ITGA1* | Integrin alpha-1 | Integrin |
| *THY1* | Thy-1 membrane glycoprotein | Cell adhesion molecule |
| **Extracellular organelle** | | |
| ***YWHAB*** | **Tyrosine 3-monooxygenase** | Scaffold/Adaptor Protein |
| ***RAC1*** | **Ras-related C3 Botulinum toxin substrate 1** | Small GTPase |
| ***RAC2*** | **Ras-related C3 Botulinum toxin substrate 2** | Small GTPase |
| ***MFGE8*** | **Lactadherin** | Oxidoreductase |
| ***HNRNPL*** | **Heterogeneous nuclear ribonucleoprotein L** | RNA processing factor |
| *BAIAP2* | Brain-specific angiogenesis inhibitor 1-associated protein 2 | Scaffold/Adaptor Protein |
| *LAMC1* | Laminin subunit gamma-1 | Extracellular matrix protein |
| *SYT4* | Synaptotagmin-4 | Membrane trafficking regulatory protein |
| *SFRP1* | Secreted frizzled-related protein 1 | Signaling receptor |
| *STXBP1* | Syntaxin-binding protein 1 | Membrane trafficking regulatory protein |
| **Vesicle** | | |
| ***YWHAB*** | **Tyrosine 3-monooxygenase** | Scaffold/Adaptor Protein |
| ***RAC1*** | **Ras-related C3 Botulinum toxin substrate 1** | Small GTPase |
| ***RAC2*** | **Ras-related C3 Botulinum toxin substrate 2** | Small GTPase |
| ***MFGE8*** | **Lactadherin** | Oxidoreductase |
| ***HNRNPL*** | **Heterogeneous nuclear ribonucleoprotein L** | RNA processing factor |
| *HGF* | Hepatocyte growth factor | Serine protease |
| *LIG3* | Leucine-rich repeats and immunoglobulin-like domains protein 3 | Transmembrane signaling receptor |
| *TMED4* | Transmembrane Emp24 Domain-containing protein 4 | Vesicle coat protein |
| *CDC42* | Cell division control protein 42 homolog | Small GTPase |
| *LAMA5* | Laminin subunit alpha-5 | Extracellular matrix protein |

**Supplementary table 2: Marker proteins identified in exosome preparations from human induced microglia**

The protein composition of exosomes purified from $SORLA^{WT}$, $SORLA^{N1358S}$, or $SORLA^{KO}$ iMG was determined by mass spectrometry. The table lists 58 identified proteins (from a total of 598 proteins) included in the top 100 list of most expressed proteins in exosomes (http://microvesicles.org/extracellular_vesicle_markers).

| Gene | Protein | Gene | Protein |
|---|---|---|---|
| *ACTB* | Actin, cytoplasmic 1 | *HSPA8* | Heat shock cognate 71 kDa protein |
| *ACTN1* | Alpha-actinin-1 | *IQGAP1* | Ras GTPase-activating-like protein |
| *ACTN4* | Alpha-actinin-4 | *KPNB1* | Importin subunit beta-1 |
| *AHCY* | Adenosyl homocysteinase | *LDHA* | L-lactate dehydrogenase A chain |
| *ANXA1* | Annexin A1 | *LDHB* | L-lactate dehydrogenase B chain |
| *ANXA11* | Annexin A11 | *MSN* | Moesin |
| *ANXA2* | Annexin A2 | *MYH9* | Myosin-9 |
| *ANXA5* | Annexin A5 | *PDCD6IP* | Programmed cell death 6-interacting protein |
| *ANXA6* | Annexin A6 | *PFN1* | Profilin-1 |
| *ANXA7* | Annexin A7 | *PGAM1* | Phosphoglycerate mutase 1 2 |
| *C3* | Complement C3 | *PGK1* | Phosphoglycerate kinase 1 |
| *CAP1* | Adenylyl cyclase-associated protein 1 | *PKM* | Pyruvate kinase |
| *CFL1* | Cofilin-1 | *PPIA* | Peptidyl-prolyl cis-trans isomerase A |
| *CLIC1* | Chloride intracellular channel protein 1 | *PRDX1* | Peroxiredoxin-1 |
| *CLTC* | Clathrin heavy chain 1 | *PRDX2* | Peroxiredoxin-2 |
| *EEF1A1* | Elongation factor 1-alpha 1 | *RAB10* | Ras-related protein Rab-10 |
| *EEF2* | Elongation factor 2 | *RAB7A* | Ras-related protein Rab-7a |
| *EHD1* | EH domain-containing protein 1 | *RAP1B* | Ras-related protein Rap-1b |
| *EIF4A1* | Eukaryotic initiation factor 4A-I | *TPI1* | Triosephosphate isomerase |
| *ENO1* | Alpha-enolase | *TUBB4B* | Tubulin beta-4B chain |
| *EZR* | Ezrin | *VCL* | Vinculin |
| *FASN* | Fatty acid synthase | *VCP* | Endoplasmic reticulum ATPase |
| *FLNA* | Filamin-A | *YWHAB* | 14-3-3 protein beta/alpha |
| *GAPDH* | Glyceraldehyde-3-phosphate dehydrogenase | *YWHAE* | 14-3-3 protein epsilon |
| *GDI2* | Rab GDP dissociation inhibitor beta | *YWHAG* | 14-3-3 protein gamma |
| *GNAI2* | Guanine nucleotide-binding protein G(i) subunit alpha-2 | *YWHAQ* | 14-3-3 protein theta |
| *GPI* | Glucose-6-phosphate isomerase | *YWHAZ* | 14-3-3 protein zeta/delta |
| *GSN* | Gelsolin | | |
| *HSP90AB1* | Heat shock protein HSP 90-beta | | |
| *HSP90AA1* | Heat shock protein HSP 90-alpha | | |
| *HSPA5* | Endoplasmic reticulum chaperone | | |

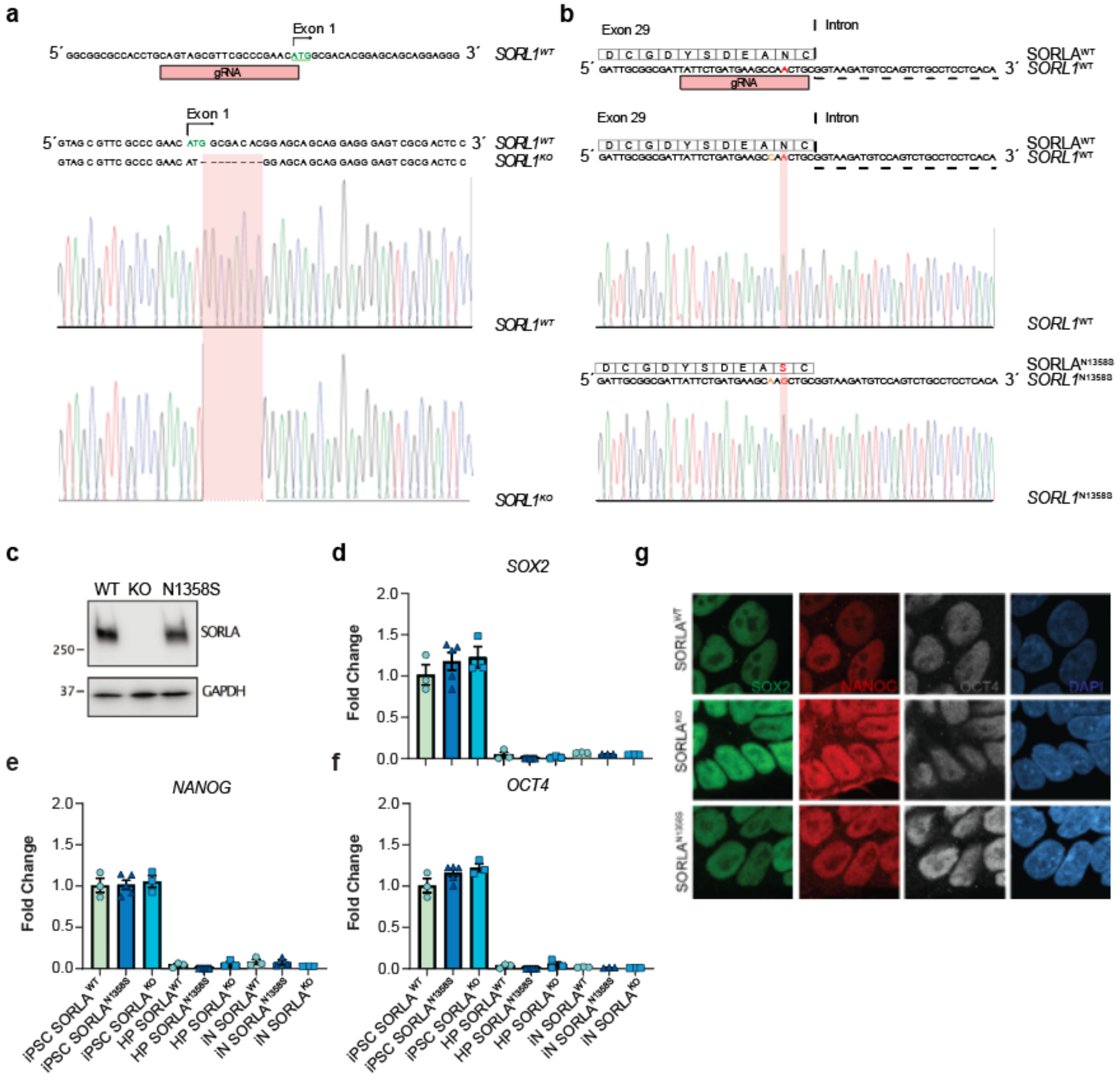


**Supplementary figure 1: Characterization of iPSC lines wildtype or genetically deficient for SORLA, or expressing SORLA$^{N1358S}$**

(**a**) CRISPR/Cas9 strategy used to disrupt *SORL1* in the human iPSC line HMGUi001-A. Placement of the guide RNA (gRNA) at the ATG start codon of the wildtype (WT) *SORL1* locus is shown. Sequence analysis of the *SORL1* locus documents an 8 nt deletion at the start codon in SORLA-deficient cells (KO) (**b**) CRISPR/Cas9 strategy used to introduce the N1358S mutation into cell line HMGUi001-A. Sequence analyses documents a change at amino acid position 1358 from asparagin (N) to serine (S) in SORLA$^{N1358S}$ iPSCs (in red). In addition, a silent mutation was introduced into the preceding codon for alanine (A; in yellow), a strategy used to improve accuracy of Cas9 cleavage (see method section for detail) (14). (**c**) Western blot analysis of SORLA in iPSCs lysates confirms ablation of the receptor in SORLA$^{KO}$, but comparable receptor levels in SORLA$^{N1358S}$ and SORLA$^{WT}$. Detection of *GAPDH* served as loading control. Migration of molecular weight markers in the gel are indicated in kDa. (**d-f**) Quantitative (q) RT-PCR analysis of transcript levels for pluripotency markers *OCT4*, *NANOG*, and *SOX2* in iPSCs of the indicated *SORL1* genotypes are shown. Levels are given as fold change compared to SORLA$^{WT}$ (set to 1). Data represent three independent differentiation experiments (n = 3-5 biological replicates). (**g**) Immunocytochemical detection of pluripotency markers OCT4 (white), NANOG (red), and SOX2 (green) in iPSC lines of the indicated *SORL1* genotypes. Nuclei are counterstained with DAPI (blue). Data are shown as individual channel configurations. Scale bar: 20 µm.

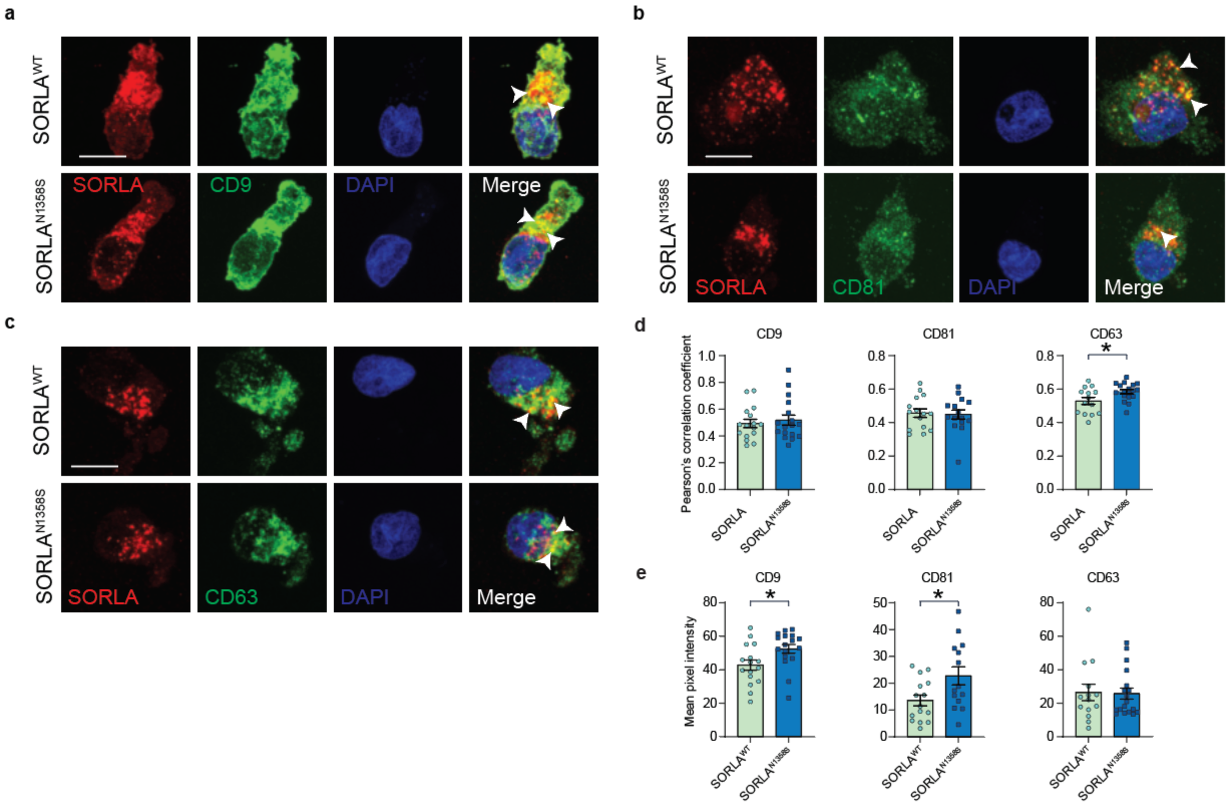


**Supplementary figure 2: Colocalization of SORLA$^{WT}$ and SORLA$^{N1358S}$ with exosomal biogenesis markers in human microglia**

(**a - c**) Immunofluorescence detection of SORLA (red) and exosomal biogenesis markers CD9 (green, a), CD81 (green, b), or CD63 (green, c) in SORLA$^{WT}$ or SORLA$^{N1358S}$ iMG. Nuclei were counterstained with DAPI (blue). Single as well as merged channel configurations are shown. White arrowheads in the merged images indicate areas of co-localization of SORLA and the respective marker. Scale bar: 10 µm. (**d, e**) Co-localization of SORLA$^{WT}$ or SORLA$^{N1358S}$ with the given exosomal biogenesis markers. The extent of co-localization was determined using Pearson´s correlation coefficient. The mean pixel intensity was based on a mask covering the full cell body (e) (n=17 - 20 cells per condition from two independent differentiation experiments). Statistical significance was determined using Student’s t-test (*, $p<0.05$).
